# Season and city shape urban bioaerosol composition beyond vegetation and socioeconomic gradients

**DOI:** 10.1101/2025.09.25.678581

**Authors:** Sarah Poirier, Jonathan Rondeau-Leclaire, Maria Faticov, Alexis Roy, Gaële Lajeunesse, Jean-François Lucier, Sarah Tardif, Steven W. Kembel, Carly Ziter, Catherine Laprise, Alain Paquette, Catherine Girard, Isabelle Laforest-Lapointe

**Affiliations:** Département de biologie, Université de Sherbrooke, Sherbrooke, Québec, Canada; Réseau de recherche en santé durable lié à la qualité de l’air et de l’environnement sonore (AIRS), CHU Sainte-Justine, Montréal, Québec, Canada; Centre d’Étude de la Forêt, Département des sciences biologiques, Université du Québec à Montréal, Montréal, Québec, Canada; Département des sciences biologiques, Université du Québec à Montréal, Montréal, Québec, Canada; Department of Wildlife, Fish, and Environmental Studies, Swedish University of Agricultural Sciences, Umeå, Sweden; Centre de calcul scientifique, Université de Sherbrooke, Sherbrooke, Québec, Canada; Department of Biology, Concordia University, Montréal, Québec, Canada; Département des Sciences Fondamentales, Université du Québec à Chicoutimi, Chicoutimi, Québec, Canada; Centre intersectoriel en santé durable, Université du Québec à Chicoutimi, Saguenay, QC, Canada; Département de biochimie, de microbiologie et de bio-informatique, Université Laval, Québec, Canada; Centre d’études nordiques (CEN), Université Laval, Québec, Québec, Canada

**Keywords:** urban ecology, bacteria, fungi, pollen, microbial ecology, metagenomics, aerobiome

## Abstract

Urban vegetation varies with socio-economic gradients, as lower-income neighborhoods often host sparser and less diverse green spaces. This disparity may affect respiratory health by influencing exposure to bioaerosols. Understanding the characteristics of this aerobiome could help anticipate risks related to allergies and other respiratory conditions. Here, we hypothesized that urban vegetation cover and socio-economic status shape urban bioaerosols. We sampled bioaerosols at 65 sites across three Canadian cities of varying population size and density using an active air sampler over four months, and characterized their bacterial, fungal, and pollen composition using amplicon sequencing. Seasonal alpha diversity varied significantly for fungi and pollen. Based on beta diversity, sampling period alone explained up to 40% of pollen, 29% of fungal, and 11% of bacterial community composition variation. In contrast, vegetation cover explained only a minor portion of the variance in bioaerosol composition, and median household income, almost none. These findings provide a critical baseline for understanding the urban aerobiome and highlight the need to study how vegetation identity and diversity, rather than cover alone, may shape bioaerosol dynamics in cities. As cities grow and urban greening initiatives expand, demystifying the aerobiome dynamics becomes an urgent public health priority.

## INTRODUCTION

Biodiversity, from microscopic bacteria to majestic trees, provides countless ecosystem services that are essential for human population health worldwide^1^. However, the benefits of biodiversity are increasingly threatened as global change drives a rise in disturbances across biomes, including higher temperatures, more frequent and intense natural disasters, and land-use change^2^. Such pressures are leading to losses of biodiversity^3^ and ecosystem functions^4,5^, threatening human health and well-being^6^. In urban areas, where 77% of Canadians^7^ and 57% of the global population^8^ currently reside, these impacts are particularly acute. Cities across the world are expanding rapidly and are projected to host over 70% of the global human population in the next 30 years^9^. Urban centers are major sources of airborne particles and gases due to dense human activity, infrastructure, and land-use change^10^. While there is significant attention on urban vegetation as a way to mitigate pollutant exposure in cities – alongside other co-benefits^1,11^ – vegetation also influences the quantity and diversity of bioaerosols^12,13^ (also known as the “aerobiome”), which in turn can influence human health^14,15^. Yet, the determinants of urban bioaerosol composition and dynamics remain poorly understood. In this work, we focus on two key components of urban bioaerosols: plant tissues and microbes (bacteria and fungi), aiming to uncover how urban environmental characteristics influence their presence and dynamics.

Bioaerosols are airborne biological particles originating from diverse sources, including soil, water bodies, plant surfaces, animal waste, and human activities such as agriculture and industry^16^. In urban areas, they commonly include bacteria, fungi, and plant tissues such as pollen^17,18^, a well-known allergen. While typically harmless, pollen can trigger allergic reactions in sensitized individuals, with symptoms varying seasonally and geographically^19^. Urban vegetation can exacerbate airborne pollen exposure, particularly if the urban canopy is dominated by dioecious male trees and anemophilous tree species that produce more pollen^18,20,21^. This suggests that the health impacts of urban vegetation extend beyond its benefits^22^, warranting a broader focus on bioaerosols in public health and urban biodiversity research. However, the current methods for predicting airborne pollen concentration in urban environments are limited by the sparse distribution of monitoring stations and the labor-intensive nature of microscopic identification, which restricts both spatial resolution and real-time responsiveness^23^.

Microorganisms represent the most abundant and diverse component of urban biodiversity, colonizing soils, plants, animals, insects, the built environment, with many taxa dispersing through the air^24^. Land-use patterns have been shown to shape airborne microbial communities or “microbiomes”^25,26^, and recent studies highlight the role of local vegetation in influencing airborne bacterial composition in both urban^27^ and natural settings^28^. Advances in high-throughput sequencing are expanding our understanding of the urban microbiome^29^. Nevertheless, most studies to date have focused on indoor environments, leaving the outdoor “aerobiome” comparatively understudied^27,30,31^. This is in part due to the difficulty of studying highly dynamic microbial communities, which are characterized by rapid evolution and high dispersal potential^32–34^. Understanding these dynamics is especially important in cities, where biodiversity, including microbial exposure, is unevenly distributed and closely tied to social inequalities^35^.

Despite its recognized benefits for health and well-being, access to urban vegetation remains inequitably distributed in many cities. Indeed, studies have shown that high-income neighborhoods tend to have more abundant and diverse tree communities than low-income ones^36–39^. This may result in large disparities in bioaerosol exposure and respiratory health outcomes. The urban aerobiome and its variation across socioeconomically diverse districts are critical, yet often-overlooked, determinants of population health. Exposure to diverse microbial and plant-derived bioaerosols, particularly during early life, has been linked to a decrease in the risk of developing asthma and allergies^40^ through immune system development^41^. Autoimmune and allergic diseases, such as asthma and atopic dermatitis, affect between 10% and 30% of the population globally, with higher prevalence observed in high-income countries and among children^42–44^. Moreover, children in urban centers face a 70% higher risk of asthma, independent of ethnicity and income^45^. To mitigate these trends, urban planning initiatives must integrate a mechanistic understanding of how vegetation influence human health through bioaerosols.

In cities, local vegetation diversity has been linked to microbial communities in key buildings (e.g., schools, hospitals, and homes)^46–48^. A recent study in Finland demonstrated that a 28-day biodiversity intervention in urban children increased both skin and gut diversity of *Gammaproteobacteria*, with positive effects on immunoregulatory pathways^49^. These findings support the idea that enhancing urban biodiversity could reduce the risk of immune-mediated diseases^49^. However, historical legacies have led to higher vegetation cover and diversity in low-risk (high income) neighborhoods, thus potentially reducing their public health potential^50^. Maintaining or enhancing vegetation cover and diversity in high-risk (low-income) neighborhoods, where respiratory illnesses are more prevalent^51^, could yield greater health gains and help reduce environmental health disparities. These socioeconomic disparities in access to urban vegetation highlight the need to consider social equity in urban greening strategies^50,52,53^. Ignoring the disparate spatial stratification of vegetation from low-income to high-income districts could limit the equity and effectiveness of urban biodiversity interventions. Therefore, understanding how vegetation cover and median income interact to shape bioaerosol exposure is essential as a first step towards designing equitable and health-promoting cities.

In this study, we assessed the composition and diversity of airborne microorganisms (bacterial and fungal) and pollen (but also additional plant debris) across three Canadian cities (Montréal, Québec City, and Sherbrooke) differing in population size and density (Fig. 1; Supplementary Fig. S1, Table S1;). Samples were collected across the summer by active air sampling at 65 sites (Fig. 1A) in Montréal (25 sites; Fig. 1B), in Québec City (25 sites; Fig. 1C), and in Sherbrooke (15 sites; Fig. 1D). For each city, sampling site coordinates were determined according to two gradients: (1) median household income data by postal code^54^ and (2) vegetation cover using the normalized difference vegetation index (NDVI)^55^. This design allowed us to identify sites along contrasting levels of both income and vegetation gradients (e.g., low vegetation cover and low socioeconomic status vs. high vegetation cover and low socioeconomic status).

**Figure 1.**
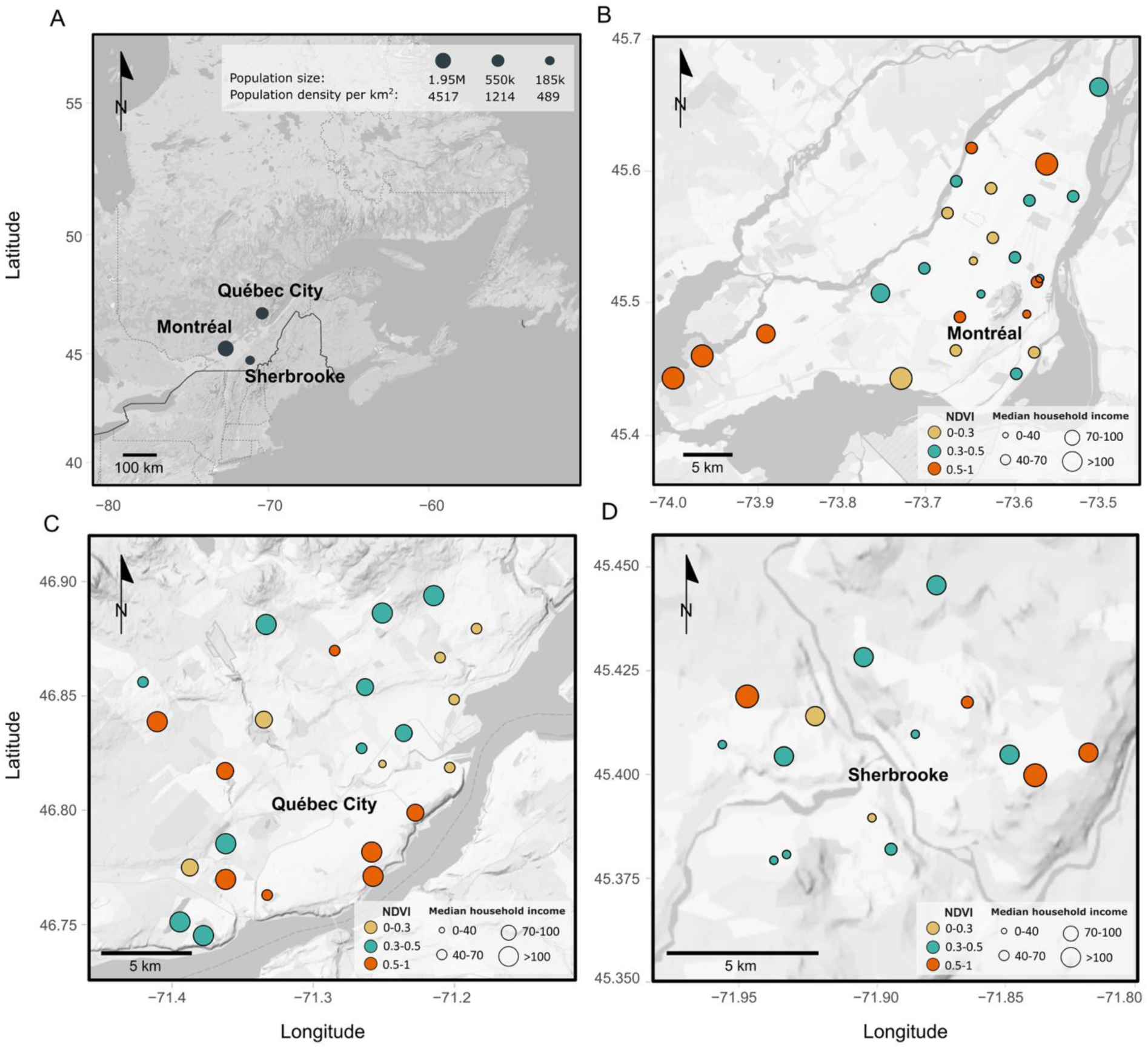
Sampling locations and their characteristics. **(A)** Geographic location of the three study cities in Québec, Canada. **(B–D)** Sampling distribution of the 25 sites within Montréal **(B)**, 25 sites within Québec City **(C)**, and 15 sites within Sherbrooke **(D)**. Circle size indicates median household income in Canadian dollars (thousands of CA$), with smaller circles representing lower-income areas. Circle color corresponds to the Normalized Difference Vegetation Index (NDVI), reflecting vegetation density. The vegetation density gradient (NDVI) is categorized in three bins (1: 0–0.3, 2: 0.3–0.5, 3: 0.5–1) and the median household income gradient is categorized in four bins (< 40k CA$, 40k–70k CA$, 70k–100k CA$, and > 100k CA$). The Montréal study design is part of the *Montreal Urban Observatory*.

Our main objective was to characterize the composition and diversity of urban bioaerosols and to estimate the relative influence of urban vegetation and socio-economic gradient on bioaerosol dynamics. Specifically, we evaluated how bacterial, fungal, and pollen composition, diversity and abundance vary in relation to (i) city identity, (ii) sampling period, (iii) vegetation cover (NDVI), and (iv) median household income. To achieve this, we employed high-throughput sequencing targeting the 16S rRNA (bacteria), ITS (fungi), and *trnL* (pollen) genes, complemented by quantitative PCR (qPCR) to estimate total bacterial load in bioaerosols.

## RESULTS

### Urban bioaerosol composition varies across season and cities

We assessed the influence of city identity, sampling period, vegetation cover, and median household income on the composition of bacterial, fungal, and pollen bioaerosols (Table 1; Fig. 2). Of note, Québec City could only be sampled in spring and fall because of limited access to the active air sampler. Based on Bray-Curtis dissimilarities, sampling period was the strongest determinant of variation in community composition of bacteria, fungi, and pollen (R² = 10.7%, 29.2%, and 39.7%, respectively), followed by city identity (R² = 7.9%, 8.6%, 7.6%,). Vegetation cover, measured via normalized difference vegetation index (NDVI)^55^, was a statistically significant but weak predictor of bioaerosol composition (R² = 1.2%, 0.5%, 0.5%,). Median household income showed a minor effect, limited to bacterial community composition (R² = 0.8%). We also found a significant interaction between city and sampling period (R² = 3.0%, 7.7%, 9.3%). Turnover-nestedness analyses indicated that compositional changes were predominantly driven by amplicon sequencing variants (ASVs, see Methods) turnover rather than shifts in the relative abundance of persistent bioaerosol taxa, a pattern consistent across bacterial, fungal, and pollen composition (Supplementary Fig. S4). The magnitude of compositional differences between cities and sampling periods was so pronounced that it produced a horseshoe effect in the PCoA ordinations (Fig. 2), a well-documented artifact in unconstrained ordination methods that typically arises when strong gradients (e.g., in Fig. 2BC) dominate the dataset^56^. This pattern motivated us to analyze bioaerosol datasets separately for each city, Montréal (Fig. 3), Québec City (Fig. 4), and Sherbrooke (Fig. 5), rather than aggregating them, allowing for clearer interpretation of local dynamics.

**Figure 2.**
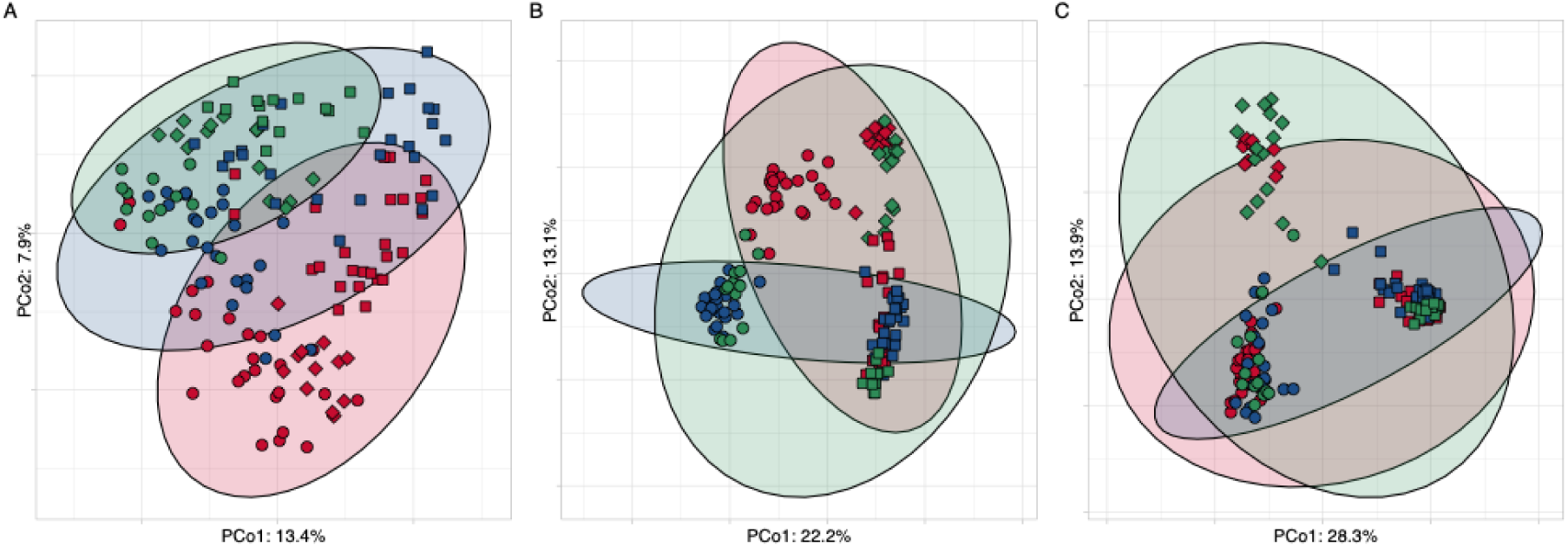
Principal Coordinates Analysis (PCoA) ordinations on Bray-Curtis dissimilarities of bioaerosol communities across three Canadian cities. Ordinations are shown for (A) bacteria, (B) fungi, and (C) pollen sampled in Montréal (red), Québec City (blue), and Sherbrooke (green). Sampling periods are represented by distinct shapes: spring (squares), summer (diamonds), and fall (circles).

**Figure 3.**
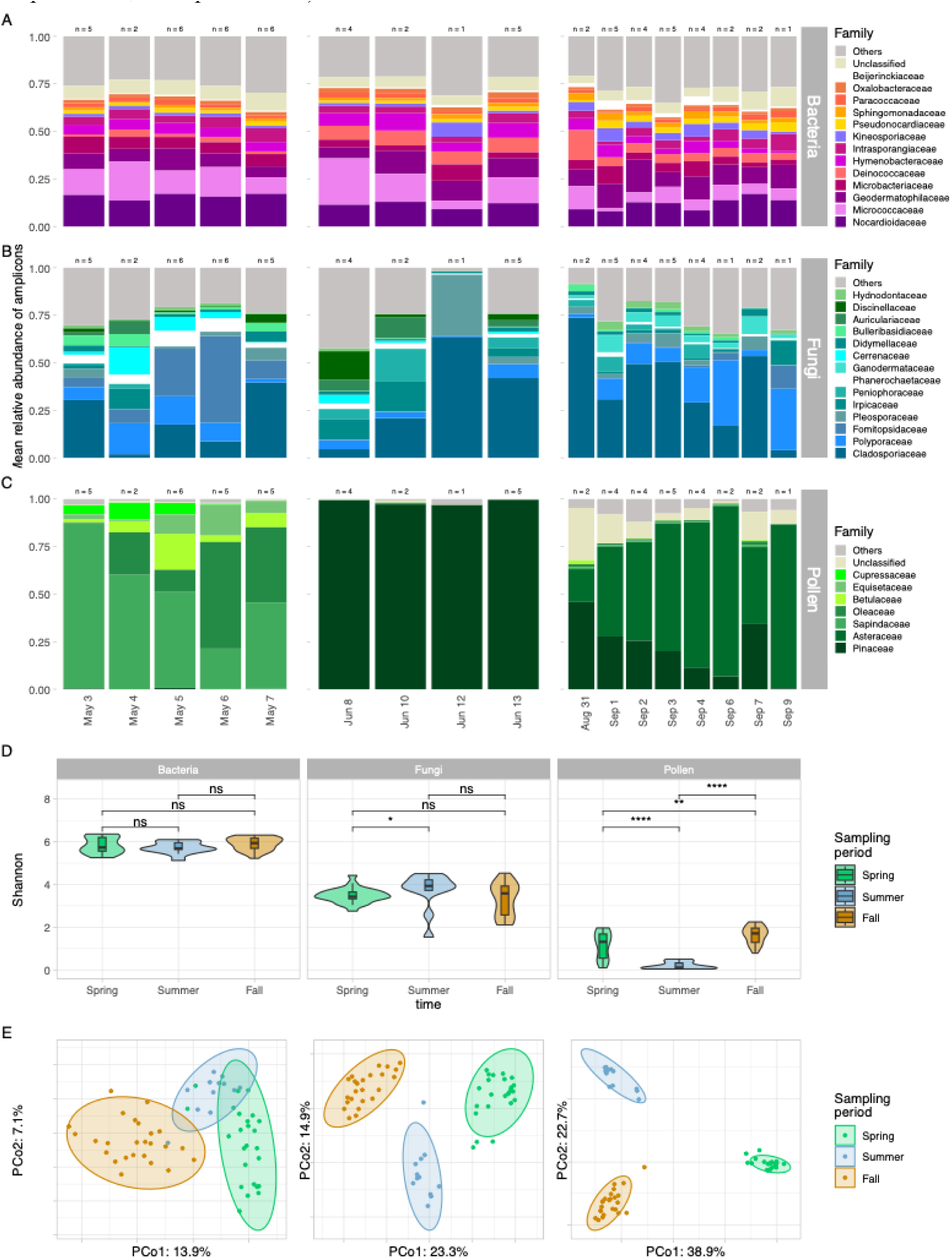
Composition and diversity of aerosols in Montréal. **(A-C)** Relative abundance of dominant families for bacteria **(A)**, fungi **(B)**, and pollen **(C)** in urban air. The barcharts are structured in three sampling periods and show the mean relative abundance by date of sampling. The n shows the number of samples aggregated per day but sequencing was performed on each sample separately. **(D)** Alpha diversity (Shannon index) across sampling periods for each biological group. **(E)** Principal coordinates analysis (PCoA) of Bray-Curtis dissimilarities showing seasonal shifts in community structure. Statistical significance was assessed using the Wilcoxon signed-rank test with p-value correction using the Holm procedure; asterisks indicate significant differences (*p < 0.05, **p < 0.01, ***p < 0.001, ****p < 0.0001).

**Figure 4.**
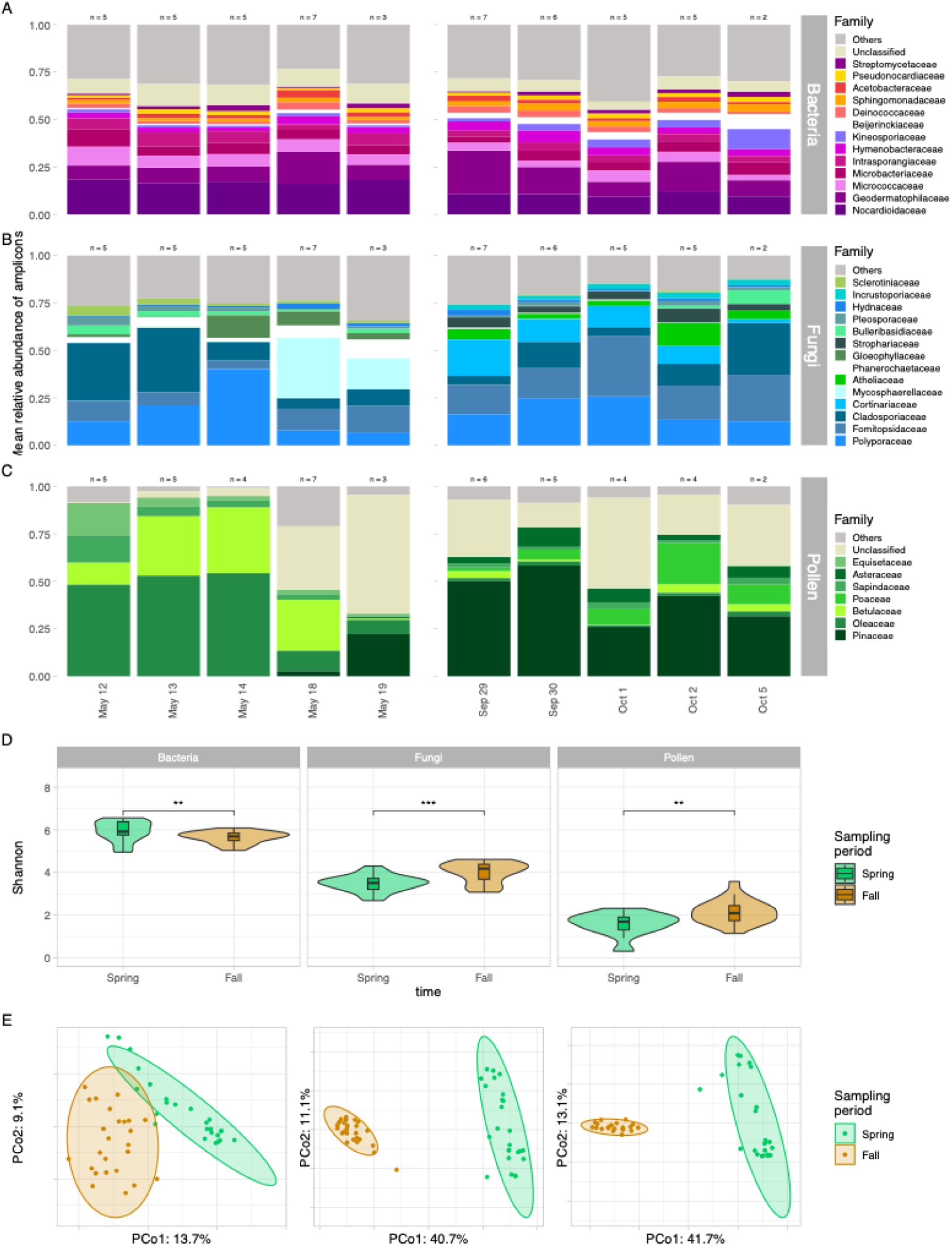
Composition and diversity of aerosols in Québec. **(A-C)** Relative abundance of dominant families for bacteria, fungi, and pollen in urban air. The barcharts are structured in three sampling periods and samples are aggregated by date of sampling. The n shows the number of samples aggregated per day. **(D)** Alpha-diversity (Shannon index) across sampling periods for each biological group. **(E)** Principal coordinates analysis (PCoA) of Bray-Curtis dissimilarities showing seasonal shifts in community structure. Statistical significance was assessed using the Wilcoxon signed-rank test with p-value correction using the Holm procedure; asterisks indicate significant differences (*p < 0.05, **p < 0.01, ***p < 0.001, ****p < 0.0001).

**Figure 5.**
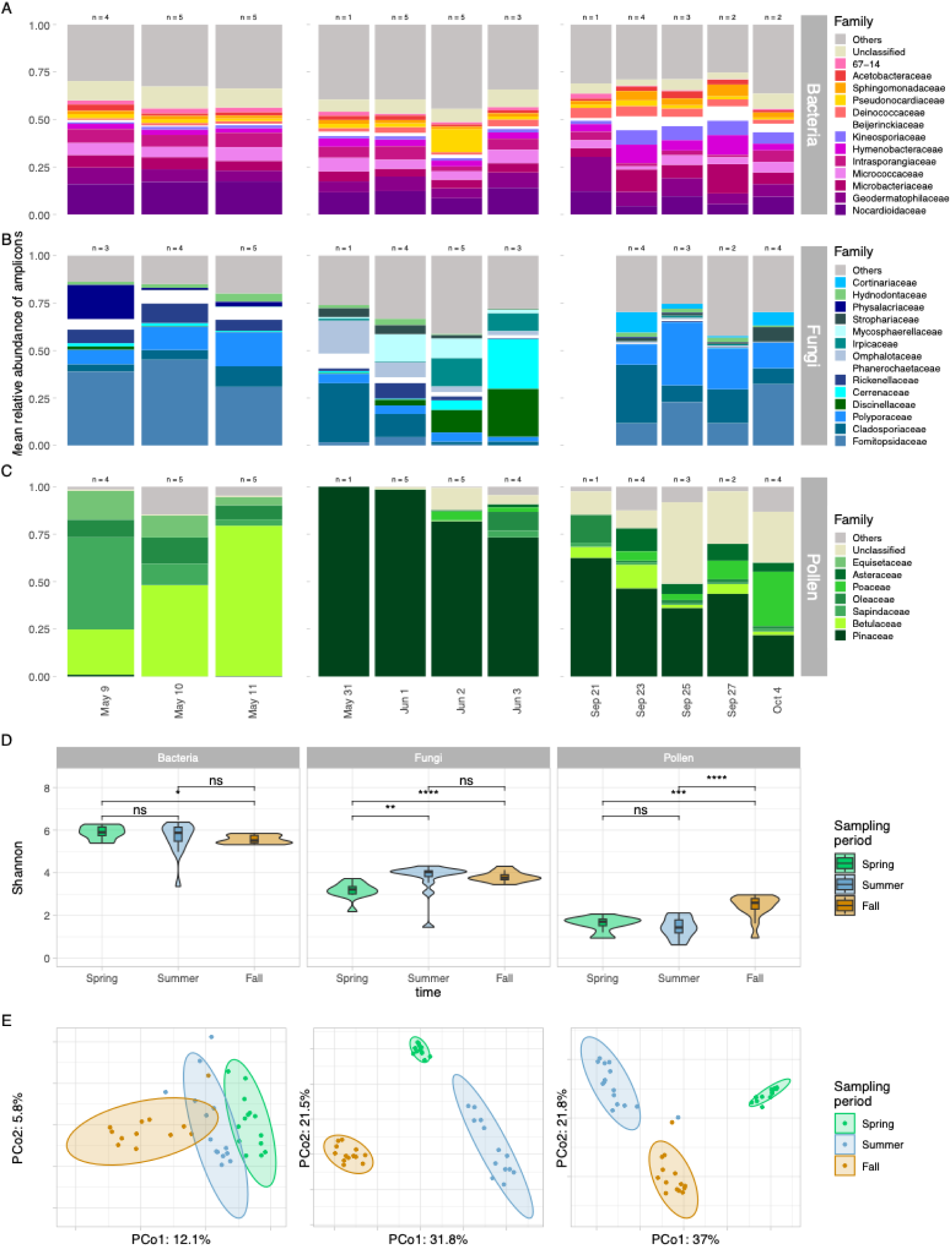
Composition and diversity of aerosols in Sherbrooke. **(A-C)** Relative abundance of dominant families for bacteria, fungi, and pollen in urban air. The barcharts are structured in three sampling periods and samples are aggregated by date of sampling. The n shows the number of samples aggregated per day. **(D)** Alpha-diversity (Shannon index) across sampling periods for each biological group. **(E)** Principal coordinates analysis (PCoA) of Bray-Curtis dissimilarities showing seasonal shifts in community structure. Statistical significance was assessed using the Wilcoxon signed-rank test with p-value correction using the Holm procedure; asterisks indicate significant differences (*p < 0.05, **p < 0.01, ***p < 0.001, ****p < 0.0001).

**Table 1.** Variation in bioaerosol community composition across cities and seasons assessed by PERMANOVA. Statistical analysis of bacterial, fungal, and pollen communities reveals significant differences in composition across urban environments (Montréal, Québec City, Sherbrooke) and sampling periods (SP; spring, summer, fall), as determined by permutational multivariate analysis of variance (PERMANOVA) on Bray-Curtis dissimilarities. *F:* pseudo-F value, DF: degrees of freedoms, R^2^: % r-squared, *p*: observed *p*-value.

| *Variables* | *Bacteria* |  |  |  | *Fungi* |  |  |  | *Pollen* |  |  |  |
| --- | --- | --- | --- | --- | --- | --- | --- | --- | --- | --- | --- | --- |
|  | *F* | *DF* | *R*<sup>2</sup> | *p* | *F* | *DF* | *R*<sup>2</sup> | *p* | *F* | *DF* | *R*<sup>2</sup> | *p* |
| City | 7.34 | 2 | 7.9 | 0.0001 | 11.12 | 2 | 8.6 | 0.0001 | 12.34 | 2 | 7.6 | 0.0001 |
| Sampling Period (SP) | 9.98 | 2 | 10.7 | 0.0001 | 37.67 | 2 | 29.2 | 0.0001 | 64.13 | 2 | 39.7 | 0.0001 |
| City:SP | 1.86 | 3 | 3.0 | 0.0002 | 6.59 | 3 | 7.7 | 0.0001 | 10.07 | 3 | 9.3 | 0.0001 |
| NDVI | 2.3 | 1 | 1.2 | 0.0006 | 1.4 | 1 | 0.5 | 0.0921 | 1.73 | 1 | 0.5 | 0.0179 |
| Income (IN) | 1.4 | 1 | 0.8 | 0.0407 | 1.08 | 1 | 0.4 | NS | 1.31 | 1 | 0.4 | NS |
| NDVI:IN | 1.11 | 1 | 0.6 | NS | 0.81 | 1 | 0.3 | NS | 0.92 | 1 | 0.3 | NS |
| **Total explained** | **24.2%** |  |  |  | **46.8%** |  |  |  | **57.9%** |  |  |  |

### Distinct taxonomic and relative abundance patterns in urban bioaerosols

Urban aerosols composition was generally more stable in bacterial communities compared to fungi and pollen (Figs. 2–4ABC; Supplementary Figs. S5–S7). Specifically, within any given sampling time and city combination, Bray-Curtis dissimilarities between bacterial ASV compositions were much lower and less variable than between fungal or plant compositions (Supplementary Fig. S8). Most bacterial sequences were assigned to two phyla: *Actinomycetota* (65.7%) and *Pseudomonadata* (15.0%) (Figs. 3A, 4A, 5A; Supplementary Fig. S5, Table S2A). To explore the concept of a core microbiome, we identified ASVs present in at least 95% of samples and with an overall mean relative abundance of at least 0.1%^57^. Using this criterion, we identified eight bacterial ASVs as core members, all of them from the *Actinobacteria* (Supplementary Table S3). These taxa included two variants from the genus *Nocardioides* (family *Nocardioidaceae*), three from the genus *Blastococcus* (family *Geodermatophilaceae*), and one unclassified member of the order *Frankiales*. Additionally, two variants from the family *Micrococcaceae*, belonging to the genera *Arthrobacter* and *Kocuria*, were also identified as core taxa.

Fungal sequences were assigned to the phyla Basidiomycota (63.0%) and Ascomycota (37.0%) (Figs. 3B, 4B, 5B; Supplementary Fig. S6, Table S2B). We identified eight fungal ASVs as core members (Supplementary Table S3). These taxa included three ASVs from the *Ascomycota* (class *Dothideomycetes*) and five from the *Basidiomycota* (class *Agaricomycetes*). Within the *Ascomycota*, two ASVs belonged to the genus *Cladosporium* (family *Cladosporiaceae*), and one to *Alternaria* (family *Pleosporaceae*), both genera are well-known contributors to airborne fungal spores^58,59^. Among the *Basidiomycota*, two ASVs were assigned to *Bjerkandera* (family Phanerochaetaceae), and one each to *Sistotremastrum* (*Hydnodontaceae*), *Trametes* (*Polyporaceae*), and *Resinicium* (*Rickenellaceae*). These five ASVs are wood-decaying fungi commonly found in forested and urban green spaces^60,61^.

The plant (pollen, but also including other aerial plant debris) component of bioaerosols (Figs. 3C, 4C, 5C; Supplementary Fig. S7, Table S2C) was dominated by the *Pinaceae* family across all cities (35.0% overall; 33.3% in Montréal, 24.8% in Québec, 47.0% in Sherbrooke), comprising conifers such as *Abies balsamea* (balsam fir), *Pinus* spp. (pines), *Picea* spp. (spruces), and *Larix laricina* (tamarack). These species typically release pollen around June, aligning with the peak observed during the second sampling time (93.3% of sequences). Notably, *Pinaceae* ASVs remained prevalent into the fall sampling period (September-October), accounting for 32.8% of sequences. The *Asteraceae* family were also strongly detected in our sequences (13.9%; 28.0% in Montréal, 3.1% in Québec, 2.9% in Sherbrooke), with a strong presence in fall in Montréal (59.6%). *Oleaceae*, which includes *Fraxinus* spp. (ashes) and *Syringa* spp. (lilacs), followed (9.8% overall; 24.0%, 34.7%, and 10.1%, respectively in spring), with peak abundance in early May. Only one ASV was identified as a core member of the pollen dataset, and it was assigned to the genus *Syringa* (family *Oleaceae*; Supplementary Table S3). The *Sapindaceae* (e.g., *Acer saccharum*, *Acer platanoides*) showed early-season peaks (9.8% overall; 52.8%, 5.2%, 21.0%, respectively in spring). The *Betulaceae*, including *Corylus* spp. (hazels), *Betula* spp. (birches), *Alnus* (alders), and *Ostrya virginiana* (hop-hornbeams), contributed significantly to spring bioaerosols (8.7% overall; 7.5%, 21.1%, and 50.4%, respectively. Finally, the *Poaceae*, a family of grasses and weedy species linked to hay fever (e.g., *Poa pratensis*, *Phleum pratense*, *Bromus inermis*, *Digitaria sanguinalis*, *Elytrigia repens*), were primarily detected in fall, and especially outside Montréal (9.5% and 9.3% in the fall for Québec and Sherbrooke).

### Contrasting patterns in alpha and beta diversity across urban bioaerosol types

Across all cities, for bacteria, diversity measured by the Shannon index remained stable across sampling periods (Supplementary Fig. S9). In contrast, fungal alpha diversity was lower during spring compared to summer (p < 0.001) and fall (p < 0.001). Pollen alpha diversity, on the other hand, showed a distinct seasonal trend, with higher diversity in the fall and lower in the summer (all pairwise comparisons: p < 0.001).

City-level comparisons (Supplementary Fig. S10) revealed occasional season-specific trends in alpha diversity. In spring, only fungal alpha diversity was found to be significantly lower in Sherbrooke compared to Montreal (p < 0.05). In summer, pollen alpha diversity was higher in Sherbrooke than in Montréal (p < 0.0001). In the fall, bacterial alpha diversity was highest in Montréal, compared to Québec (p < 0.01) and Sherbrooke (p < 0.01). Montréal exhibited lower fungal alpha diversity than Québec (p < 0.01), and lower pollen alpha diversity than both Québec City (p < 0.01) and Sherbrooke (p < 0.001).

Temporal trends within each city further highlighted these dynamics. In Montréal (Fig. 3D), pollen alpha diversity was drastically lower in summer compared to spring and fall (both p < 0.0001). In Québec (Fig. 4D), bacterial alpha diversity was higher in spring (p < 0.01), while both fungal and pollen diversity were higher in fall (p < 0.001 and p < 0.01, respectively). In Sherbrooke (Fig. 5D), fungal alpha diversity was higher in summer (p < 0.01) and fall (p < 0.0001) compared to spring, while pollen diversity peaked in fall compared to spring (p < 0.001) and summer (p < 0.0001). Finally, no significant differences were found in alpha diversity across vegetation index category (NDVI) or median household income (Supplementary Figs. S9–S13).

For beta diversity, principal coordinates analysis (PCoA) revealed a distinct clustering of samples by sampling period for all three amplicons studied (Fig. 3E, 4E, 5E). In Montreal, the first two axes captured 21%, 38.2% and 61.6% of inertia in Bray-Curtis dissimilarities for bacteria, fungi and pollen, respectively. In Québec, they captured 22.8%, 51.8% and 55.1%, respectively, while in Sherbrooke, 17.9%, 53.3% and 58.8%, respectively.

### Differential abundance of bioaerosol taxa

We performed differential abundance analyses using ANCOM-BC2 to identify temporal shifts in absolute abundances of bioaerosol genera, setting summer as the reference season and controlling for city identity (Fig. 6; effect sizes and p-values listed in Supplementary Table S7). Genera were identified as differentially abundant if they met 3 conditions: enriched or depleted in at least one season compared to summer; p_adj._ < 0.01; and passed ANCOM-BC2’s sensitivity test (see Methods).

**Figure 6.**
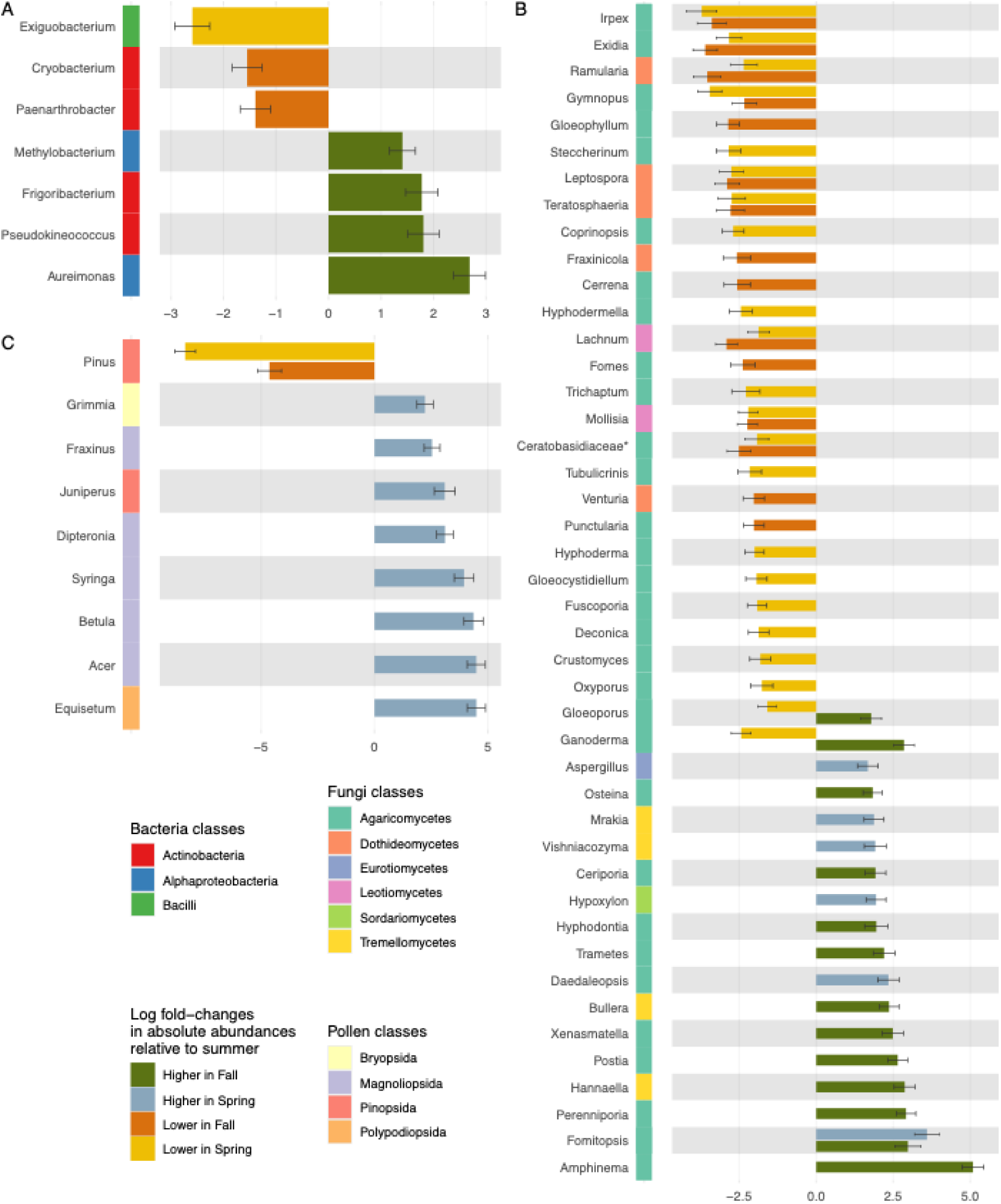
Differentially abundant genera across seasons identified by ANCOM-BC. Differential abundance analysis of bioaerosols using ANCOM-BC2 Dunnett tests, with summer as the reference season and city identity included as a covariate. Positive values indicate enrichment in spring (blue) or fall (green) relative to summer, while negative values indicate depletion in spring (yellow) or fall (orange). Error bars represent standard errors of the estimated log-fold (LF) changes in absolute abundance. **(A)** Bacterial genera; **(B)** plant (pollen) genera; **(C)** fungal genera. Only genera with at least 20% prevalence across samples were tested, and only those with at least one significant comparison having passed the pseudocount sensitivity test, and with an absolute LF-change >1.5 for fungi are shown. Significance threshold is q < 0.01, where q is the p-value adjusted for multiple testing using the Holm procedure. _*genus *incertae sedis*_.

Among bacterial genera (Fig. 6A), *Exiguobacterium* showed significantly lower abundance in spring, while *Cryobacterium* and *Paenarthrobacter* showed lower abundance in fall. Fall samples were characterized by increased abundance of *Frigobacterium*, *Pseudokineococcus*, *Aureimonas*, and *Methylobacterium* sequences. Interestingly, all *Dothiodeomycetes* and *Leotiomycetes* identified as differentially abundant were enriched in summer compared to fall and often spring too. Similarly, all *Tremelloycetes* identified by this analysis were depleted in the summer relative to either spring or fall. In contrast, no consistent patterns of enrichment were observed for the *Agaricomycetes* (Fig. 6A).

Multiple fungal genera were identified as differentially abundant, revealing distinct temporal patterns in urban air (Fig. 6B). Several genera, including *Irpex*, *Exidia*, and *Ramularia*, were significantly enriched in summer samples, suggesting a peak in fungal sporulation or dispersal during warmer months. In contrast, spring samples were enriched in genera such as *Daedaleopsis* and *Aspergillus*, many of which include cold-tolerant or early-season taxa. Multiple genera were enriched in fall, including *Amphinema*, *Perenniporia, Postia*, *Trametes*, and *Ceriporia*, suggesting a shift in urban fungal community composition as temperatures decline. Some genera showed reduced abundance in specific seasons. For example, *Gloeophyllum*, *Fraxinicola*, and *Venturia* were less abundant in fall compared to summer, while *Steccherinum*, *Coprinopsis*, and *Hyphodermella*, were less abundant in spring. Interestingly, the genus *Formitopsis* was more abundant in both spring and fall compared to summer, suggesting a bimodal seasonal pattern. In addition, the genera *Ganoderma* and *Gloeoporus* showed significant depletion in spring compared to summer, followed by an enrichment in fall, suggesting a gradual seasonal increase (Fig. 6B).

Finally, differential abundance analysis of pollen composition also demonstrated strong temporal variation (Fig. 6C). *Pinus* was significantly more abundant in summer compared to both spring and fall, consistent with its expected mid-season pollination peak. In contrast, several taxa, including *Fraxinus*, *Juniperus*, *Dipteronia*, and *Equisetum*, were enriched in spring, suggesting early-season pollen release. Additional spring-associated genera included *Grimmia*, *Syringa*, *Betula*, and *Acer*, many of which are common in urban landscapes and contribute substantially to early-season airborne pollen loads (Fig. 6C).

### Bacterial bioaerosol quantity varies between cities and to lesser extent across seasons

In terms of quantity estimates, the qPCR analysis showed that bacteria were more abundant in summer than fall in Montréal (*p adj.* = 0.028) Fig. 7A). There were no significant differences in Québec or Sherbrooke across sampling periods. When comparing cities, our results show that there were less bacterial bioaerosols in Sherbrooke than in Montréal or Québec City at each sampling periods (Fig. 7B). None of the other gradients (median income and vegetation) could be associated with airborne bacteria quantity (Fig. S14).

**Figure 7.**
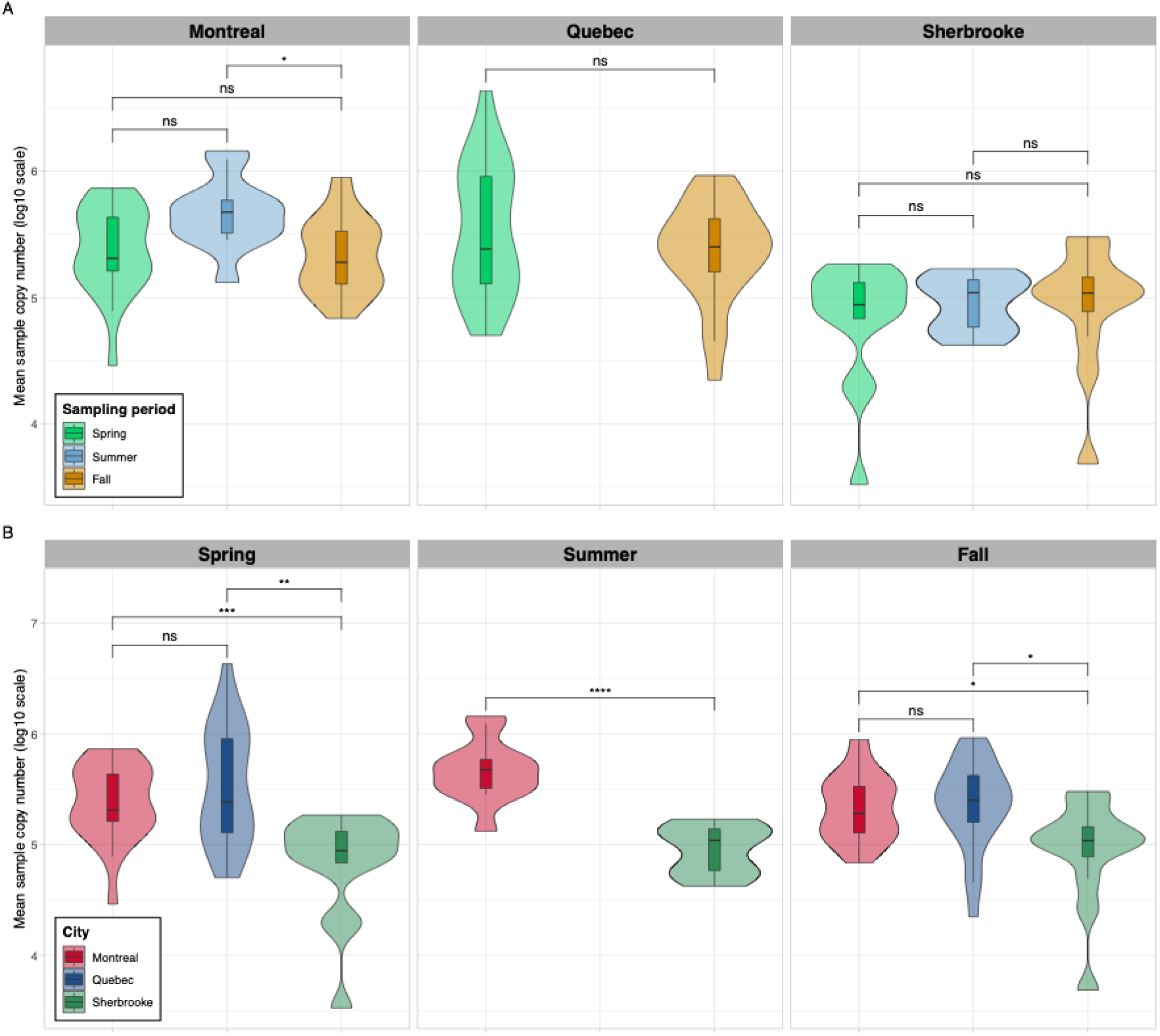
Abundance of bacterial urban bioaerosols across cities and sampling periods. Extreme values with copy number > 5e+06 were excluded from the statistical test and plots. Values shown are log10(mean copy number). Mean copy number is the mean of three qPCR replicates per sample. Statistical significance established with pairwise Wilcoxon test with p-value correction using the Holm procedure. Statistical significance was assessed using the Wilcoxon signed-rank test with p-value correction using the Holm procedure; asterisks indicate significant differences (*p < 0.05, **p < 0.01, ***p < 0.001, ****p < 0.0001).

## DISCUSSION

In this study, we investigated the composition, diversity, and abundance of urban bioaerosols, focussing on bacteria, fungi, and pollen (including plant debris), across three Canadian cities varying in size and population density (Fig. 1). Our central objective was to disentangle the relative influence of environmental and socio-economic drivers, including vegetation cover, neighborhood socio-economic status, and sampling period, on bioaerosol dynamics. While vegetation is known to shape bioaerosol abundance and composition^12,13^ and often correlates with socio-economic gradients^36–39^, the interplay between these factors and the urban aerobiome remains poorly understood. Our findings revealed that sampling period was the most influential driver of community composition across all bioaerosol types, explaining 10.7–39.7% of the observed variation (Table 1). City identity and its interaction with sampling period also showed significant effects, accounting for 7.6–8.6% and 3.0–9.3% of the variation, respectively (Table 1). In contrast, vegetation cover and socio-economic status appeared to have limited influence on bioaerosol community structure at this spatial resolution. These results underscore the importance of temporal dynamics in shaping urban bioaerosols, revealing that temporal variation far outweighs local vegetation or socio-economic gradients in driving microbial and pollen composition. This insight challenges prevailing assumptions that urban vegetation and neighborhood characteristics are dominant determinants of airborne biodiversity and instead points to the need for time-sensitive monitoring and policy interventions. Understanding when bioaerosols shift most dramatically can inform public health strategies, urban planning, and climate resilience efforts, especially as cities face increasing pressures from environmental change and population growth.

In line with previous studies, our results confirm that sampling period is the dominant driver of variation in bioaerosols, including bacteria, fungi, and pollen, across urban environments^62–65^ (Table 1, Fig. 2–5). This strong temporal signal likely reflects seasonal shifts in abiotic conditions such as temperature, humidity, rainfall, and plant phenology that influence the growth, release, and aerosolization of biological particles^66^. Additionally, microbial succession on plant surfaces (the phyllosphere) varies seasonally, contributing to fluctuations in airborne microbial communities^13,67^. The second most influential factor was city identity, which may reflect differences in vegetation identity^68^, topography^69^, land use^26^, road traffic^70^, pollution^71^, and climate^72^. For instance, higher traffic volumes and distinct climatic conditions in Montréal compared to Québec City and Sherbrooke (Table S4) may contribute to divergent bioaerosol dynamics. While previous studies have reported that urban vegetation influences airborne microbial communities^12,13^, our results suggest only a minor effect of vegetation cover on bioaerosol composition. This discrepancy may stem from differences in spatial resolution or from the fact that earlier studies often inferred vegetation influence based on the presence of plant-associated taxa in the air, rather than direct measurements of surrounding vegetation^73^. These findings highlight the primacy of time and place in shaping the urban aerobiome, suggesting that when and where we sample matters more than how green or affluent a neighborhood is. This has important implications for urban health monitoring, allergen forecasting, and environmental justice, as it calls for dynamic, city-specific strategies rather than static, one-size-fits-all approaches to managing bioaerosol exposure in cities.

The most relatively abundant bacterial families detected across all seasons and cities included the *Nocardioidaceae*, *Geodermatophilaceae*, *Micrococcaceae*, and *Microbacteriaceae* (Figs. 3A, 4A, 5A). All belong to the phylum *Actinobacteriota*, class *Actinobacteria*, which includes many taxa commonly found in soil and aquatic environments^74^, both recognized as major sources of airborne microorganisms^75^. Notably, *Nocardioidaceae* has been previously identified in bioaerosols^75^, and its members are known for their resilience in harsh conditions. These Gram-positive bacteria can survive long-range atmospheric transport, including via sandstorms originating from desert topsoils^76^, and have also been detected in urban air samples^77^. Our results also showed limited variation in alpha diversity across seasons and cities, with only a slight decrease from spring to fall in Québec (Fig. 4), and a modest increase in Montréal compared to Québec and Sherbrooke during the fall (Supplementary Fig. S10). The consistent presence of *Actinobacteria* across diverse urban environments and seasons suggests a stable core of airborne bacterial taxa that may be shaped more by regional environmental reservoirs rather than by local urban features. This stability at the ASV level, coupled with the resilience of taxa such as *Nocardioidaceae*, highlights the potential for long-distance microbial dispersal and raises important questions about the ecological roles and health implications of persistent airborne bacteria in cities.

Interestingly, our results show that *Cryobacterium*, a genus of obligately psychrophilic bacteria that has been sampled from glaciers in Antarctica^78,79^, is more abundant in summer compared to fall (Fig. 6). Additionally, *Methylobacterium* (Fig. 6), known to be present in air, soil, and on plant leaves, where some of them can produce plant growth promoting substances^80^, was most abundant in the fall. This could be caused by falling and decaying of leaves, during which leaf bacteria could get aerosolized in the air. This genus is also part of the *Beijerinckiaceae* family, which contains numerous plant-colonizing taxa. These findings reveal how temporal ecological processes, such as plant growth and decay, shape the composition of airborne microbial communities in cities. The differential depletion of *Cryobacterium* in fall and enrichment of *Methylobacterium* in fall highlights the dynamic link between terrestrial ecosystems and urban air, suggesting that bioaerosol composition reflects not only environmental conditions but also biological activity on the ground. This has implications for understanding seasonal exposure risks, urban ecosystem connectivity, and the potential for airborne microbes to influence plant and human health, particularly as climate change alters phenological patterns, microbial dispersal dynamics, and the timing of allergen peaks in urban environments.

The identification of a small, consistent set of core ASVs across diverse urban environments suggests the presence of a resilient airborne bacterial community that persists regardless of local conditions (Table S3). The dominance of *Actinobacteria*, many of which are known for their environmental robustness, points to a shared microbial signature in urban air, potentially shaped by common sources such as soil, dust, and plant surfaces. Recognizing these core taxa is essential for understanding baseline urban bioaerosol composition, which can inform future studies on air quality and microbial exposure.

Although genomic sequencing offers a powerful means of characterizing the composition and diversity of bioaerosols, it remains limited in its ability to quantify absolute abundances. To address this limitation, we employed quantitative PCR (qPCR) to estimate bacterial bioaerosol concentrations. Our results indicate a temporal trend, with bacterial abundance peaking in summer and declining in fall (Fig. 7A). This pattern contrasts with previous studies reporting lower airborne bacterial loads during summer months^81^. Among the three cities studied, Sherbrooke consistently exhibited the lowest concentrations of airborne bacteria (Fig. 7B), a finding that diverges from observations by Xie et al., who reported comparable bacterial bioaerosol levels across large urban centers^82^. One possible explanation for Sherbrooke’s lower levels is its urban morphology: the relative absence of tall buildings may reduce the entrapment of airborne particles, thereby enhancing air circulation and dispersal of bioaerosols. These findings highlight the importance of accounting for local urban characteristics in bioaerosol studies and open new avenues for research into how urban morphology and atmospheric dynamics shape the spatiotemporal distribution of airborne microbial communities.

Three fungal families dominated the sample sequences across all cities: *Cladosporiaceae* (21.6%) *Polyporaceae* (13.2%), and *Fomitopsidaceae* (12.1%). In comparison with bacteria (R^2^ = 10.7%), sampling period explained 29.2% of the variation in fungal community structure (Table 1). In addition to this increase in compositional similarity within season, fungal alpha diversity was lower in spring compared to summer and fall in Québec and Sherbrooke, but not in Montréal (Figs. 3–5). This increase in fungal alpha diversity in fall was also previously observed by Núñez et al.^165^. The identification of a fungal core microbiome composed of ubiquitous allergenic taxa (e.g., *Cladosporium*, *Alternaria*)^58,83^ and wood-decaying fungi highlights the diverse ecological origins of airborne fungi in cities (Table S3). These findings suggest that urban air consistently reflects inputs from vegetation, soil, and decaying organic matter. Understanding the composition and stability of fungal bioaerosols is crucial to assess seasonal allergen exposure, ecosystem connectivity, and the potential impacts of urban planning on airborne fungal diversity.

For pollen, composition was primarily driven by sampling period (Table 1; Figs. 2– 5), consistent with previous studies^84,85^. These temporal and city-specific patterns likely reflect differences in local vegetation profiles, shaped by urban canopy composition and flowering phenology. Daily fluctuations in pollen composition (Fig. 5) suggest that site-specific environmental factors, such as fine particles, temperature, and humidity, may further influence airborne pollen dynamics. These differences between cities could stem from variations in tree species composition, urban forestry practices, and microclimatic conditions, all of which contribute to the timing and intensity of pollen release. This work provides a critical baseline for understanding how urban vegetation and seasonal dynamics shape airborne pollen exposure. As cities continue to grow and climate conditions shift, such insights are essential to anticipate changes in allergen loads and informing urban planning and public health strategies aimed at mitigating respiratory health risks. For instance, city-specific pollen profiles and their seasonal dynamics could inform the development of targeted public health advisories or the strategic placement of air filtration systems in schools, hospitals, and elderly care facilities during peak allergen periods.

While this study offers novel insights into the temporal dynamics of urban bioaerosols, several limitations should be considered when interpreting the results. Due to logistical constraints related to access to the active air sampler, we were unable to sample sites in Québec City during the summer, which may have limited our ability to fully capture seasonal variation. Additionally, although we initially planned to conduct shotgun metagenomic sequencing to quantify microbial communities, the quality and quantity of extracted DNA was insufficient. Future studies could explore alternative sampling materials or preservation methods to improve DNA integrity, as well as extend sampling durations to better capture temporal variability. It is also important to note that short-read amplicon sequencing detects DNA from both viable and non-viable organisms, meaning that the presence of a taxon does not necessarily indicate biological activity. Furthermore, the restricted combinations of socio-economic and vegetation gradients may have reduced our power to detect subtle effects. As such, the absence of statistically significant results should not be interpreted as evidence of no effect. Although we did not find strong evidence of differential exposure to bioaerosols across socio-economic gradients, this study provides a valuable foundation for understanding how urban infrastructure and environmental context shape airborne microbial communities in cities. Finally, we did not account for the potential influence of fine particulate matter (e.g., PM2.5), which may interact with bioaerosols and modulate their respiratory health impacts, including by enhancing the allergenicity of pollen.

## CONCLUSION

In this study, we show that the composition of urban bioaerosols, including bacteria, fungi, and pollen, is strongly influenced by sampling period and city identity, while alpha diversity remained relatively stable across these factors. We did not detect strong effects of vegetation cover or median household income, suggesting that socio-economic status may not substantially alter exposure to diverse bioaerosols in the urban air, though further research is needed to reach consensus. As global change accelerates, shifts in temperature, humidity, and the frequency of extreme climatic events such as widespread fires are likely to influence bioaerosol dynamics. Future studies should explore the role of vegetation identity and diversity, which may better explain the presence of plant-associated microbial taxa in the air than vegetation cover (NDVI) alone. Given the rising prevalence of respiratory conditions such as allergies and asthma, it is crucial to understand the sources and ecological drivers of urban bioaerosols, including contributions from vegetation and human activity, to better anticipate and mitigate their impacts on public health.

## METHODS

### Study design

The cities were chosen to represent a gradient of population size and density, and topography (Supplementary Table S1). We performed active air sampling to capture seasonal variation in urban bioaerosols (Supplementary Figs. S2–S3). Montréal is one of the most populated cities in Canada (population of ∼1,760,000; density of 4,500 inhabitants/km^2^; 430 km^2^), making it an important urban center to include in the study as it represents high-density urbanism and facilitates potential comparisons with cities of similar densities across the world. Montréal has the lowest cover of vegetation between the three cities with 69.1% (0.5827 normalized difference vegetation index [NDVI]) of vegetation^55^, and its topography is the one with the least variation and the lowest average elevation at 30 m^86^ (Table S1). The Montréal study design is part of the “*Montreal Urban Observatory*”, a research platform which aims to monitor urban forest ecosystems for global change adaptation and health (Paquette *et al.* in review). In addition to considering vegetation cover and household income, the Montréal study design also accounted for population density. Québec City is a medium size city (population of ∼550,000; 1,230 inhabitants/km^2^; 485 km^2^), offers a contrasting vegetation cover with 85.9% (0.6518 NDVI) of vegetation^55^, and displays the highest variation in topography (Table S1) with an average altitude of 117 m^87^. Finally, Sherbrooke is a sparsely populated city (population of ∼170,000; 490 inhabitants/km^2^; 370 km^2^) with the highest vegetation cover of 90.6% (0.6837 NDVI)^55^ and a varied topography (average altitude of 232 m^88^; Table S1), thus representative of smaller urban areas that are surrounded by agricultural fields and forests.

Within each city, to characterize environmental and socioeconomic gradients, we extracted normalized difference vegetation index (NDVI) values from Landsat-8 satellite imagery (USGS, 2021) at 2 m spatial resolution using Google Earth Engine. For each sampling site, we calculated the mean NDVI within a 200 m radius buffer to capture local vegetation cover. Household income data were obtained from the 2016 Canadian Census (Statistics Canada, 2017), which reports median total household income for 2015. Each sampling site was assigned the median household income of its corresponding Dissemination Area (DA), the smallest standardized geographic unit in Canada (population 400–700). Thus, each site was characterized by one NDVI value and one household income value, which were used as continuous predictors in subsequent models.

### Bioaerosol sampling

At each site, samples were collected by concentrating urban air on electret filters (electrostatically charged) with an aerosol collector (SASS®4100, Research International) which collects particles with an aerodynamic diameter ranging from 0.5 to 10 µm^89^. Air was sampled for one hour at 4000 L/min, 1 m aboveground ^90^. The samples were taken over three sampling rushes: from May 2^nd^ to May 19^th^, 2022 (65 samples: 25 in Montréal, 25 in Québec, and 15 in Sherbrooke), from May 31^st^ to June 13^th^, 2022 (30 samples: 15 in Montréal and 15 in Sherbrooke), and from August 31^st^ to October 5^th^, 2022 (65 samples: 25 in Montréal, 25 in Québec, and 15 in Sherbrooke). Climate measurements for temperature, relative humidity, and wind speed were taken with an enviro-meter^TM^ (Fisherbrand). Public daily recordings of temperature and precipitation^91^ are showed in Supplementary Figs. S2–S3 with dates of sampling in each city overlayed. Sampling was performed as much as possible 24h after the last precipitation, a strategy that had to be adjusted sometimes due to timeline constraints (Fig. S3). Public recordings of monthly abiotic conditions^91^ are also showed in Supplementary Table S4.

### DNA extraction

Electret filters were submitted to a treatment prior to DNA extraction modified from Mbareche *et al.*^92^. Filters were cut to remove the plastic ring with a sterilised scalpel inside a sterile biological hood. Filter membranes were transferred to a 50 mL sterile plastic tube with 10 mL of extraction buffer (1.9 mM of Na_2_H_2_PO_4_*2H_2_O, 8.1 mM of Na_2_HPO_4_, 138 mM of sodium chloride, 2.7 mM of potassium chloride, 15.4 mM of sodium azide and 0,05% (w/v) of Triton X-100®) and vigorously agitated using a Vortex for 10 min. Filter membranes were then squeezed to remove the excess liquid with sterile pliers. The extracted buffer was transferred to a Amicon® Ultra-15 Centrifugal Filter Unit from Millipore Sigma (UFC910096) and then centrifuged at 4000g for 2 min. The resulting 100 µL was then transferred to the first tube of the QIAGEN DNeasy PowerSoil Pro Kit (47016) to extract DNA, following the manufacturer’s instructions with a mean yield of 1.75 ng/µl. Controls were added to confirm that samples were not contaminated during sampling, cutting away the filter membranes from the plastic ring and extracting DNA.

### Quantitative analysis

Following DNA extraction, quantitative real-time PCR (qPCR) was performed to determine the absolute abundance of bacteria and fungi in all samples. All reactions were conducted using a Bio-Rad CFX96 system with CFX Maestro software (v2.3). Each 10 µL reaction contained 1× SYBR™ Green, 10 µM of each primer, and 2 µL of sample DNA. For bacterial quantification, primers 799F (forward) and 1193R (reverse) were used. The amplification protocol consisted of an initial denaturation at 95°C for 10 min, followed by 40 cycles of 10 s at 95°C, 10 s at 55°C, and 10 s at 95°C, with a final extension at 72°C for 10 s. Melting curve analysis was performed at the end of each run to confirm primer specificity. All reactions were conducted in triplicate. Absolute bacterial abundance was calculated by converting Cq values to *16S* rRNA gene copy numbers using standard curves generated from serial dilutions of *Erwinia amylovora* ATCC15854 DNA. The known *16S* gene copy number was derived from genome size and DNA concentration. Standard curves were included on each qPCR plate to minimize amplification bias.

### Sequencing

For bacteria, the *16S* rRNA gene V5-V6 region was amplified using the chloroplast-excluding 799F and 1115R primers^93^. For fungi, the primers ITS1F^94^ and ITS2^95^ were used to amplify the ITS1 region of the *ITS* gene. For pollen, the c-A49325 and d-B49863 primers were used to amplify the *trnL* (UAA) intron gene^96^. The resulting DNA was submitted to marker gene short read sequencing (Illumina Miseq PE300; see primers in Table S5). Raw sequences contained 35,895 ± 12,817 (16S), 33,424 ± 15,106 (ITS) and 24,900 ± 13,839 (*trnL*) reads, respectively.

### Sample composition estimation

Amplicon sequencing variants (ASVs) were resolved with the DADA2 1.26.0^97^ pipeline using R 4.4.0^98^ on high performance computer clusters hosted by the Digital Research Alliance of Canada. Amplicon sequence set (16S, ITS and *trnL* samples) were processed separately to create three distinct datasets. Briefly, primers were removed using cutadapt 2.10^99^, thus ensuring read-through reverse complements were removed. Reads with a maximum expected error of 2 or at least one base with a quality of Q = 2 were removed. 16S sequences were truncated at lengths of 230 bases (forward) and 120 bases (reverse), dropping sequences shorter than these thresholds, while the variable-length *ITS* and *trnL* sequences were dropped if they were shorter than 100 bases. Of note, after discussion with the DADA2 authors, it was determined that the highly variable length of the *trnL* would better resolve ASVs by using only the forward reads. Extensive testing of various read merging strategies on this dataset confirmed that this was the best way to prevent artificial inflating of ASV diversity while optimizing their taxonomic assignment (see the following discussion with the authors: https://github.com/benjjneb/dada2/issues/2091). Therefore, for the *trnL* dataset only, forward sequences were truncated at 275 bp and any shorter reads were discarded. For all dataset, the dada2 error model was used to resolve ASVs and infer sample composition using the “pseudo-pooling” method, and PCR chimera were removed using the “consensus” method.

### ASV taxonomic annotations

ASV were taxonomically labeled using the naïve bayesian classifier^100^ implemented in DADA2. Assignment of 16S and ITS ASV taxonomic labels was done using SILVA v138.1^101^ and UNITE for fungi v10.0^102^. For *trnL* ASV taxonomic assignment, a custom database was generated using an in-house pipeline. Briefly, the complete NCBI core_nt database^103^ was downloaded on May 21, 2025, comprising 115,113,413 genomic sequences from all domains of life. These sequences were indexed by Dicey (0.3.3)^104^, a bio-informatic tool that finds sequences that can be amplified by a given pair of primers. This tool was applied to the *trnL* primer pair, and the resulting set of sequence was formatted as a database to be used as reference with DADA2’s taxonomic classifier.

### ASV post-processing

ASV abundance matrix, taxonomic labels and sample metadata were consolidated and filtered independently for each amplicon dataset to create a phyloseq (1.48.0)^105^ object that served as input for all statistical analyses, ensuring referential integrity between ASVs, taxonomy, and samples. First, ASVs without a Phylum annotation or at least a sum of 50 sequences across all samples were dropped. Then, samples with fewer than 9,000 (16S) and 2,500 (ITS and *trnL*) sequences were discarded. These thresholds were established visually using sequence count distribution, aiming to increase the smallest sample count of each dataset while discarding the smallest number of samples. The resulting number of sequences, ASVs, samples, as well as the number of sequences and ASVs per sample and ASV prevalence are summarized in Table S6 and taxonomic assignation in Fig. S15 for each dataset. Given recent research, we manually changed all instances of the genus *Methylorubrum* for *Methylobacterium*^106^.

Plots were generated using ggplot2 (3.5.1), patchwork (1.3.0), cowplot (1.1.3), gtable (0.3.6), gridExtra (2,3) and MetBrewer (0.2.0).

### Diversity analyses

Diversity analyses were performed on rarefied data to account for uneven sequencing depth. Each amplicon dataset was rarefied to the number of sequences of its smallest sample, using the *rarefy_even_depth* function from the phyloseq package. Alpha diversity was computed using the Shannon index. Differences in alpha diversity across sampling periods were tested using a pairwise Wilcoxon test and *p*-values were adjusted using the Holm procedure. PCoAs were computed on rarefied data using the variance-stabilized Bray-Curtis dissimilarities between samples, which were computed using the *varianceStabilizingTransformation* function from the DESeq package and the vegdist function from the vegan (2.6.8)^108^ package, respectively. PERMANOVA were performed using the *adonis2* function of the vegan package, with 9999 permutations. For the global test across all cities, the following model evaluated by terms:

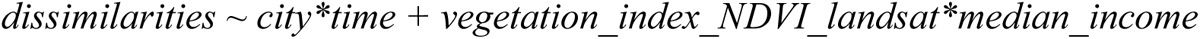

Because PERMANOVA does not separate effects of species turnover from changes in species richness, the *beta.multi* function from the *betapart* package^109^ was used to disentangle the contributions of species turnover and nestedness to total community dissimilarity. Analyses were conducted at the ASV level for 16S (bacteria), ITS (fungi), and *trnL* (pollen). Bacterial, fungal, and pollen datasets were split by city and season pair (spring–summer, summer–fall, spring–fall), retaining only replicates present in both sampling periods. For each subset, sample counts were aggregated by period and converted the data to presence/absence format using the *decostand* function from the *vegan* package. Pairwise Sørensen dissimilarities between periods were computed using *beta.multi(index.family = “sorensen”)*. This approach partitions total beta diversity (β_SOR_) into two additive components: the turnover component (β_SIM,_ where existing taxa are replaced by others) representing species replacement between periods, and the nestedness component (β_SNE_) representing changes in species richness (e.g., species losses or gains).

### Differential abundance analysis

Changes in absolute abundances were estimated using the log-linear model implemented in ANCOM-BC2 (2.4.0)^110^ on non-rarefied data and are expressed in (natural) log-fold changes. This approach was designed to account for both sample- and taxon-specific biases. It was chosen because amplicon-based sequencing is known to introduce sequencing bias, resulting in the overrepresentation of certain taxa relative to their true relative abundances. This bias was observed based on a control sample added to the *trnL* sequencing run, which was made of a known amount of DNA from a set of 17 pollen vouchers (Supplementary Figure S16), justifying the choice of a statistical method that explicitly accounts for taxon-specific biases.

The ANCOM-BC2’s Dunnett test was used to test pairwise differences in genera found in at least 20 % of samples, using summer as a reference group. Fold-changes (FC) are expressed on the natural log scale, showing relative enrichment (or depletion) in absolute abundance relative to summer. *P*-values were corrected using the Holm procedure. As the *ancombc2* function was executed once per amplicon dataset (Bacteria, Fungi, and Pollen), *p*-value correction was performed manually to account for multiple testing across these datasets, a total of 1172 tests. Only genera showing p < 0.01 were considered as differentially abundant. Additionally, ANCOM-BC2 uses pseudocounts for log ratios and performs tests to evaluate the sensitivity of inference to the pseudocount value; comparisons who did not pass this test were excluded. Considering the large number of differentially abundant taxa (especially fungi), only FCs with absolute values >1 (1.5 for fungi) are reported.

### qPCR analysis

A copy number calculator^111^ was used for each sample, with the length of the gene sequence for 16S, which was 1550 bp^112^. The average number of copies of the gene 16S (7 copies) was used, which ranged from 1 to 15 copies per organism^113^. Differences in bacterial load across sampling periods and across cities were tested using a pairwise Wilcoxon test and *p*-values were adjusted using the Holm procedure.

## AUTHOR CONTRIBUTION STATEMENT

S.P. conceived the study with I.L.L., A.P., and C.G. S.P. led the writing of the first version of the manuscript and contributed to methodology, formal analysis, investigation, visualization, data curation, and both original draft and review/editing stages. J.R.-L. contributed to formal analysis, visualization, data curation, and writing – review and editing. M.F. contributed to methodology, formal analysis, visualization, and writing – review and editing. A.R. contributed to formal analysis and writing – review and editing. G.L. contributed to methodology, formal analysis, and writing – review and editing. J.-F.L. developed software. S.T. contributed to methodology and writing – review and editing. C.L. contributed to conceptualization, funding acquisition, resources, and writing – review and editing. S.W.K. contributed to conceptualization, funding acquisition, and writing – review and editing. C.G. contributed to conceptualization, methodology, funding acquisition, resources, supervision, and both original draft and review/editing stages. A.P. contributed to conceptualization, methodology, funding acquisition, supervision, resources, validation, and writing – review and editing. I.L.-L. led project administration and supervision, as well as contributed to conceptualization, methodology, funding acquisition, supervision, resources, investigation, validation, visualization, data curation, and both original draft and review/editing stages.

## DATA AVAILABILITY

All the raw sequences have been deposited on ENA under accession number PRJEB96751. All scripts used to generate the results of this study from the raw sequences have been deposited on Github (https://github.com/jorondo1/urban_bioaerosols). These files will be rendered public upon publication.

## Supporting information

see Supporting Information

## ACKNOWLEDGEMENTS

We are grateful to the members of the Paquette laboratory at UQÀM who worked on setting up the project, as well as to Ema Lussier, Jérémy Fraysse, Joey Chamard, Sarah Ishak, Karina Gisèle Mac Si Hone, Sophie Boutin, Ève Lebeau, Jennifer Fontaine and the Duchaine laboratory at the Institut universitaire en cardiologie et pneumologie de Québec (Université Laval) for their great help in field and laboratory work. We also acknowledge the support of Calcul Québec, Compute Canada (CCS), and the Digital Research Alliance of Canada for providing the high-performance computing resources essential to our data processing and analysis.

## References

1. Sandifer, P. A., Sutton-Grier, A. E. & Ward, B. P. Exploring connections among nature, biodiversity, ecosystem services, and human health and well-being: Opportunities to enhance health and biodiversity conservation. Ecosyst. Serv. 12, 1–15 (2015).

2. Potapov, P., et al. Unprecedentedly high global forest disturbance due to fire in 2023 and 2024. Proc. Natl. Acad. Sci. 122, e2505418122 (2025).

3. Ripple, W. J. et al. World Scientists’ Warning to Humanity: A Second Notice. BioScience 67, 1026–1028 (2017).

4. Barnosky, A. D. et al. Has the Earth’s sixth mass extinction already arrived? Nature 471, 51–57 (2011).

5. Isbell, F., Tilman, D., Polasky, S. & Loreau, M. The biodiversity-dependent ecosystem service debt. Ecol. Lett. 18, 119–134 (2015).

6. Pascual, U. et al. Valuing nature’s contributions to people: the IPBES approach. Curr. Opin. Environ. Sustain. 26–27, 7–16 (2017).

7. Government of Canada, S. C. The Daily — Canada’s population estimates: Subprovincial areas, 2024. https://www150.statcan.gc.ca/n1/daily-quotidien/250116/dq250116b-eng.htm (2025).

8. World Bank Open Data. World Bank Open Data https://data.worldbank.org.

9. World Health Organization. World Health Statistics 2016: Monitoring Health for the SDGs, Sustainable Development Goals. (World Health Organization, 2016).

10. Saxena, V. Water Quality, Air Pollution, and Climate Change: Investigating the Environmental Impacts of Industrialization and Urbanization. Water. Air. Soil Pollut. 236, 73 (2025).

11. Bousquet, J. et al. Allergic rhinitis. Nat. Rev. Dis. Primer 6, 1–17 (2020).

12. Lu, C. et al. Aboveground plants determine the exchange of pathogens within air-phyllosphere-soil continuum in urban greenspaces. J. Hazard. Mater. 465, 133149 (2024).

13. Lymperopoulou, D. S., Adams, R. I. & Lindow, S. E. Contribution of Vegetation to the Microbial Composition of Nearby Outdoor Air. Appl. Environ. Microbiol. 82, 3822–3833 (2016).

14. Flies, E. J., Clarke, L. J., Brook, B. W. & Jones, P. Urbanisation reduces the abundance and diversity of airborne microbes - but what does that mean for our health? A systematic review. Sci. Total Environ. 738, 140337 (2020).

15. Rodó, X. et al. Microbial richness and air chemistry in aerosols above the PBL confirm 2,000-km long-distance transport of potential human pathogens. Proc. Natl. Acad. Sci. 121, e2404191121 (2024).

16. Lovasi, G. S. et al. Urban Tree Canopy and Asthma, Wheeze, Rhinitis, and Allergic Sensitization to Tree Pollen in a New York City Birth Cohort. Environ. Health Perspect. 121, 494–500 (2013).

17. Lai, Y. & Kontokosta, C. E. The impact of urban street tree species on air quality and respiratory illness: A spatial analysis of large-scale, high-resolution urban data. Health Place 56, 80–87 (2019).

18. Eisenman, T. S. et al. Urban trees, air quality, and asthma: An interdisciplinary review. Landsc. Urban Plan. 187, 47–59 (2019).

19. État des connaissances sur le pollen et les allergies: les assises pour une gestion efficace. (Direction de la santé environnementale et de la toxicologie, Institut national de santé publique Québec, Montréal, 2013).

20. van Dorn, A. Urban planning and respiratory health. Lancet Respir. Med. 5, 781–782 (2017).

21. Dales, R. E., Cakmak, S., Judek, S. & Coates, F. Tree pollen and hospitalization for asthma in urban Canada. Int. Arch. Allergy Immunol. 146, 241–247 (2008).

22. Roman, L. A. et al. Beyond ‘trees are good’: Disservices, management costs, and tradeoffs in urban forestry. Ambio 50, 615–630 (2021).

23. Buters, J. T. M. et al. Pollen and spore monitoring in the world. Clin. Transl. Allergy 8, 9 (2018).

24. King, G. M. Urban microbiomes and urban ecology: How do microbes in the built environment affect human sustainability in cities? J. Microbiol. 52, 721–728 (2014).

25. Burrows, S. M., Elbert, W., Lawrence, M. G. & Pöschl, U. Bacteria in the global atmosphere – Part 1: Review and synthesis of literature data for different ecosystems. *Atmospheric Chem*. Phys. 9, 9263–9280 (2009).

26. Bowers, R. M., McLetchie, S., Knight, R. & Fierer, N. Spatial variability in airborne bacterial communities across land-use types and their relationship to the bacterial communities of potential source environments. ISME J. 5, 601–612 (2011).

27. Mhuireach, G. et al. Urban greenness influences airborne bacterial community composition. Sci. Total Environ. 571, 680–687 (2016).

28. Lymperopoulou, D. S., Adams, R. I. & Lindow, S. E. Contribution of Vegetation to the Microbial Composition of Nearby Outdoor Air. Appl. Environ. Microbiol. 82, 3822–3833 (2016).

29. King, G. M. Urban microbiomes and urban ecology: How do microbes in the built environment affect human sustainability in cities? J. Microbiol. 52, 721–728 (2014).

30. Tischer, C. et al. Urban Dust Microbiome: Impact on Later Atopy and Wheezing. Environ. Health Perspect. 124, 1919–1923 (2016).

31. Afshinnekoo, E. et al. Geospatial Resolution of Human and Bacterial Diversity with City-Scale Metagenomics. Cell Syst. 1, 72–87 (2015).

32. Finlay, B. J. Global Dispersal of Free-Living Microbial Eukaryote Species. Science 296, 1061–1063 (2002).

33. Finlay, B. J. & Clarke, K. J. Ubiquitous dispersal of microbial species. Nature 400, 828–828 (1999).

34. Evans, S., Martiny, J. B. H. & Allison, S. D. Effects of dispersal and selection on stochastic assembly in microbial communities. ISME J. 11, 176–185 (2017).

35. Beninde, J., Veith, M. & Hochkirch, A. Biodiversity in cities needs space: a meta-analysis of factors determining intra-urban biodiversity variation. Ecol. Lett. 18, 581–592 (2015).

36. Tooke, T. R., Klinkenberg, B. & Coops, N. C. A geographical approach to identifying vegetation-related environmental equity in canadian cities. Environ. Plan. B Plan. Des. 37, 1040–1056 (2010).

37. Roman, L. A. et al. Human and biophysical legacies shape contemporary urban forests: A literature synthesis. Urban For. Urban Green. 31, 157–168 (2018).

38. Schell, C. J. et al. The ecological and evolutionary consequences of systemic racism in urban environments. Science 369, eaay4497 (2020).

39. Pinault, L., Christidis, T., Toyib, O. & Crouse, D. L. Ethnocultural and socioeconomic disparities in exposure to residential greenness within urban Canada. Health Rep. 32, 3–14 (2021).

40. Hanski, I. et al. Environmental biodiversity, human microbiota, and allergy are interrelated. Proc. Natl. Acad. Sci. 109, 8334–8339 (2012).

41. Valkonen, M. et al. Bacterial Exposures and Associations with Atopy and Asthma in Children. PLOS ONE 10, e0131594 (2015).

42. Savouré, M. et al. Worldwide prevalence of rhinitis in adults: A review of definitions and temporal evolution. Clin. Transl. Allergy 12, e12130 (2022).

43. Edwards-Salmon, S. E., Padmanabhan, S. L., Kuruvilla, M. & Levy, J. M. Increasing Prevalence of Allergic Disease and Its Impact on Current Practice. Curr. Otorhinolaryngol. Rep. 10, 278–284 (2022).

44. Zheng, J. et al. Global, regional, and national epidemiology of allergic diseases in children from 1990 to 2021: findings from the Global Burden of Disease Study 2021. BMC Pulm. Med. 25, 54 (2025).

45. Claudio, L., Stingone, J. & Godbold, J. Prevalence of Childhood Asthma in Urban Communities: The Impact of Ethnicity and Income. Ann. Epidemiol. 16, 332–340 (2006).

46. Kembel, S. W. et al. Architectural design influences the diversity and structure of the built environment microbiome. ISME J. 6, 1469–1479 (2012).

47. Meadow, J. F. et al. Indoor airborne bacterial communities are influenced by ventilation, occupancy, and outdoor air source. Indoor Air 24, 41–48 (2014).

48. Meadow, J. F. et al. Bacterial communities on classroom surfaces vary with human contact. Microbiome 2, 7 (2014).

49. Roslund, M. I. et al. Biodiversity intervention enhances immune regulation and health-associated commensal microbiota among daycare children. Sci. Adv. 6, eaba2578 (2020).

50. Jenerette, G. D., Harlan, S. L., Stefanov, W. L. & Martin, C. A. Ecosystem services and urban heat riskscape moderation: water, green spaces, and social inequality in Phoenix, USA. Ecol. Appl. 21, 2637–2651 (2011).

51. Gaffney, A. W., Himmelstein, D. U., Christiani, D. C. & Woolhandler, S. Socioeconomic Inequality in Respiratory Health in the US From 1959 to 2018. JAMA Intern. Med. 181, 968 (2021).

52. Gerrish, E. & Watkins, S. L. The relationship between urban forests and income: A meta-analysis. Landsc. Urban Plan. 170, 293–308 (2018).

53. Watkins, S. L. & Gerrish, E. The relationship between urban forests and race: A meta-analysis. J. Environ. Manage. 209, 152–168 (2018).

54. Gouvernement du Canada, S. C. Profil du recensement, Recensement de la population de 2021. https://www12.statcan.gc.ca/census-recensement/2021/dp-pd/prof/index.cfm?Lang=F (2022).

55. Gouvernement du Canada, S. C. Verdure urbaine et Indice de végétation par différence normalisée selon le centre de population de 2021. https://www150.statcan.gc.ca/t1/tbl1/fr/tv.action?pid=3810015801 (2022).

56. Morton, J. T. et al. Uncovering the Horseshoe Effect in Microbial Analyses. mSystems 2, e00166–16 (2017).

57. Shade, A. & Handelsman, J. Beyond the Venn diagram: the hunt for a core microbiome. Environ. Microbiol. 14, 4–12 (2012).

58. Latge, J.-P. & Paris, S. Allergens Of Alternaria And Cladosporium. in Fungal Antigens (eds Drouhet, E., Cole, G. T., De Repentigny, L., Latgé, J.-P. & Dupont, B.) 237–258 (Springer US, Boston, MA, 1988). doi:10.1007/978-1-4613-0773-0_30.

59. Kasprzyk, I. et al. Allergenic fungal spores in the air of urban parks. Aerobiologia 37, 39–51 (2021).

60. Hınçal, S. & Yalçın, M. Biological control of some wood-decay fungi with antagonistic fungi. Biodegradation 34, 597–607 (2023).

61. He, X. et al. Wood-inhabiting fungal community characteristics responses to nutrient additions vary among tree taxonomic groups. J. Soils Sediments 25, 1497– 1513 (2025).

62. Franzetti, A., Gandolfi, I., Gaspari, E., Ambrosini, R. & Bestetti, G. Seasonal variability of bacteria in fine and coarse urban air particulate matter. Appl. Microbiol. Biotechnol. 90, 745–753 (2011).

63. Bertolini, V. et al. Temporal variability and effect of environmental variables on airborne bacterial communities in an urban area of Northern Italy. Appl. Microbiol. Biotechnol. 97, 6561–6570 (2013).

64. Cáliz, J., Triadó-Margarit, X., Camarero, L. & Casamayor, E. O. A long-term survey unveils strong seasonal patterns in the airborne microbiome coupled to general and regional atmospheric circulations. Proc. Natl. Acad. Sci. 115, 12229–12234 (2018).

65. Núñez, A., García, A. M., Moreno, D. A. & Guantes, R. Seasonal changes dominate long-term variability of the urban air microbiome across space and time. Environ. Int. 150, 106423 (2021).

66. Joung, Y. S., Ge, Z. & Buie, C. R. Bioaerosol generation by raindrops on soil. Nat. Commun. 8, 14668 (2017).

67. Mhuireach, G. et al. Urban greenness influences airborne bacterial community composition. Sci. Total Environ. 571, 680–687 (2016).

68. Martin, A. J. F. et al. A biogeographical analysis of taxonomic diversity and native species dominance in 32 Canadian street tree populations. Urban Ecosyst. 28, 158 (2025).

69. Maki, T. et al. Vertical distribution of airborne bacterial communities in an Asian-dust downwind area, Noto Peninsula. Atmos. Environ. 119, 282–293 (2015).

70. Fang, Z., Ouyang, Z., Zheng, H., Wang, X. & Hu, L. Culturable airborne bacteria in outdoor environments in Beijing, China. Microb. Ecol. 54, 487–496 (2007).

71. Roy, S. & Gupta Bhattacharya, S. Airborne fungal spore concentration in an industrial township: distribution and relation with meteorological parameters. Aerobiologia 36, 575–587 (2020).

72. Polymenakou, P. N. Atmosphere: a source of pathogenic or beneficial microbes? Atmosphere 3, 87–102 (2012).

73. Lymperopoulou, D. S., Adams, R. I. & Lindow, S. E. Contribution of Vegetation to the Microbial Composition of Nearby Outdoor Air. Appl. Environ. Microbiol. 82, 3822–3833 (2016).

74. Ghai, R. et al. Metagenomics of the water column in the pristine upper course of the amazon river. PLoS ONE 6, e23785 (2011).

75. Nesterenko et al. Nocardioidaceae. John Wiley Sons Inc Bergey’s Man. Trust (2012) doi:10.1002/9781118960608.fbm00042.

76. Polymenakou, P. N., Mandalakis, M., Stephanou, E. G. & Tselepides, A. Particle size distribution of airborne microorganisms and pathogens during an intense african dust event in the eastern mediterranean. Environ. Health Perspect. 116, 292– 296 (2008).

77. Calderón-Ezquerro, M. del C., Serrano-Silva, N. & Brunner-Mendoza, C. Aerobiological study of bacterial and fungal community composition in the atmosphere of Mexico City throughout an annual cycle. Environ. Pollut. 278, 116858 (2021).

78. Suzuki, K.-I., Sasaki, J., Uramoto, M., Nakase, T. & Komagata, K. Cryobacterium psychrophilum gen. nov., sp. nov., nom. rev., comb. nov., an obligately psychrophilic actinomycete to accommodate “Curtobacterium psychrophilum” Inoue and Komagata 1976. Int. J. Syst. Evol. Microbiol. 47, 474–478 (1997).

79. Teoh, C. P., González-Aravena, M., Lavin, P. & Wong, C. M. V. L. Cold adaptation and response genes of Antarctic Cryobacterium sp. SO2 from the Fildes Peninsula, King George Island. Polar Biol. 47, 135–156 (2024).

80. Green, P. N. & Ardley, J. K. Review of the genus Methylobacterium and closely related organisms: a proposal that some Methylobacterium species be reclassified into a new genus, Methylorubrum gen. nov. Int. J. Syst. Evol. Microbiol. 68, 2727– 2748 (2018).

81. Chatoutsidou, S. E. et al. Variations, seasonal shifts and ambient conditions affecting airborne microorganisms and particles at a southeastern Mediterranean site. Sci. Total Environ. 892, 164797 (2023).

82. Xie, J., Jin, L., Luo, X., Zhao, Z. & Li, X. Seasonal disparities in airborne bacteria and associated antibiotic resistance genes in PM2.5 between urban and rural sites. Environ. Sci. Technol. Lett. 5, 74–79 (2018).

83. Kustrzeba-Wójcicka, I., Siwak, E., Terlecki, G., Wolańczyk-Mędrala, A. & Mędrala, W. Alternaria alternata and Its Allergens: a Comprehensive Review. Clin. Rev. Allergy Immunol. 47, 354–365 (2014).

84. Levac, E. A pollen calendar for the main allergenic pollen types in the borough of Lennoxville (Sherbrooke), Quebec. J. East. Townsh. Stud. 37, 43–62 (2011).

85. Lo, F., Bitz, C. M., Battisti, D. S. & Hess, J. J. Pollen calendars and maps of allergenic pollen in North America. Aerobiologia 35, 613–633 (2019).

86. Topographic maps, M. Montreal topographic map, elevation, terrain. Topographic maps https://en-ca.topographic-map.com/map-mrcm14/Montreal/?center=45.57977%2C-73.57615 (2024).

87. Topographic maps, Q. Quebec topographic map, elevation, terrain. Topographic maps https://en-ca.topographic-map.com/map-tkl5k/Quebec/ (2024).

88. Topographic maps, S. Sherbrooke topographic map, elevation, terrain. Topographic maps https://en-ca.topographic-map.com/map-k11tj/Sherbrooke/?center=45.38114%2C-71.94809 (2024).

89. Parker, A. et al. Review of Field Sampling Technologies for Characterizing Bioaerosols in Compact Spaces. (2020).

90. Provencher, J. et al. Microbial antibiotic resistance genes across an anthropogenic gradient in a Canadian High Arctic watershed. Rev. Sustain. Microbiol. (2024).

91. Canada, E. et C. climatique. Résultats de station - Données historiques - Climat - Environnement et Changement climatique Canada. https://climat.meteo.gc.ca/historical_data/search_historic_data_stations_f.html?searchType=stnProx&timeframe=1&txtRadius=25&selCity=&optProxType=park&selPark=48%7C8%7C69%7C44%7Cparc+national+Saguenay-Saint-Laurent&txtCentralLatDeg=&txtCentralLatMin=&txtCentralLatSec=&txtCentralLongDeg=&txtCentralLongMin=&txtCentralLongSec=&txtLatDecDeg=&txtLongDecDeg=&StartYear=1840&EndYear=2022&optLimit=specDate&Year=2022&Month=4&Day=5&selRowPerPage=25 (2022).

92. Mbareche, H., Veillette, M., Pilote, J., Létourneau, V. & Duchaine, C. Bioaerosols play a major role in the nasopharyngeal microbiota content in agricultural environment. Int. J. Environ. Res. Public. Health 16, 1375 (2019).

93. Aydogan, E. L., Moser, G., Müller, C., Kämpfer, P. & Glaeser, S. P. Long-term warming shifts the composition of bacterial communities in the phyllosphere of Galium album in a permanent grassland field-experiment. Front. Microbiol. 9, 144 (2018).

94. Gardes, M. & Bruns, T. D. ITS primers with enhanced specificity for basidiomycetes - application to the identification of mycorrhizae and rusts. Mol. Ecol. Mol. Genet. J. Wiley Online Libr. 2, 113–118 (1993).

95. White, T. J., Bruns, T. D., Lee, S. B. & Taylor, J. W. Amplification and direct sequencing of fungal ribosomal RNA genes for phylogenetics. in PCR - Protocols and Applications - A Laboratory Manual 315–322 (Academic Press, 1990).

96. Taberlet, P. et al. Power and limitations of the chloroplast trn L (UAA) intron for plant DNA barcoding. Nucleic Acids Res. 35, e14 (2007).

97. Callahan, B. J. et al. DADA2: High-resolution sample inference from Illumina amplicon data. Nat. Methods 13, 581–583 (2016).

98. R Core Team. R: A Language and Environment for Statistical Computing. R Foundation for Statistical Computing (2021).

99. Martin, M. Cutadapt removes adapter sequences from high-throughput sequencing reads. EMBnet.journal 17, 10 (2011).

100. Wang, Q., Garrity, G. M., Tiedje, J. M. & Cole, J. R. Naïve Bayesian Classifier for Rapid Assignment of rRNA Sequences into the New Bacterial Taxonomy. Appl. Environ. Microbiol. 73, 5261–5267 (2007).

101. Quast, C. et al. The SILVA ribosomal RNA gene database project: improved data processing and web-based tools. Nucleic Acids Res. 41, D590–D596 (2013).

102. Abarenkov, K. et al. UNITE general FASTA release for Fungi. 10.15156/BIO/2959332 (2024).

103. Sayers, E. W. et al. Database resources of the National Center for Biotechnology Information in 2025. Nucleic Acids Res. 53, D20–D29 (2025).

104. Rausch, T., Fritz, M. H.-Y., Untergasser, A. & Benes, V. Tracy: basecalling, alignment, assembly and deconvolution of sanger chromatogram trace files. BMC Genomics 21, 230 (2020).

105. McMurdie, P. J. & Holmes, S. phyloseq: an R package for reproducible interactive analysis and graphics of microbiome census data. PloS One 8, e61217 (2013).

106. Green, P. N. & Ardley, J. K. Review of the genus Methylobacterium and closely related organisms: a proposal that some Methylobacterium species be reclassified into a new genus, Methylorubrum gen. nov. Int. J. Syst. Evol. Microbiol. 68, 2727– 2748 (2018).

107. Kassambara, A. rstatix: Pipe-Friendly Framework for Basic Statistical Tests. R package version 0.7.2. https://rpkgs.datanovia.com/rstatix/index.html (2023).

108. Oksanen, J. et al. vegan: community ecology package. (2022).

109. Baselga, A. & Orme, C. D. L. betapart: an R package for the study of beta diversity. Methods Ecol. Evol. 3, 808–812 (2012).

110. Lin, H. & Peddada, S. D. Analysis of compositions of microbiomes with bias correction. Nat. Commun. 11, 3514 (2020).

111. Prediger, E. Copy Number Calculator: Convert from nanograms to copy number | IDT. Integrated DNA Technologies https://www.idtdna.com/pages/education/decoded/article/calculations-converting-from-nanograms-to-copy-number (2023).

112. Clarridge, J. E. Impact of 16S rRNA Gene Sequence Analysis for Identification of Bacteria on Clinical Microbiology and Infectious Diseases. Clin. Microbiol. Rev. 17, 840–862 (2004).

113. Lee, Z. M.-P., Bussema, C., III & Schmidt, T. M. rrnDB: documenting the number of rRNA and tRNA genes in bacteria and archaea. Nucleic Acids Res. 37, D489– D493 (2009).

