## Supplementary material for "Season and city shape urban bioaerosol composition beyond vegetation and socioeconomic gradients": see Supporting Information

**Table S1.** Characteristics for Montréal, Québec City, and Sherbrooke.

| **City** | | **Montréal**[1] | **Québec**[2] | **Sherbrooke**[3] |
|---|---|---|---|---|
| Size (km$^2$) | | 430 | 485 | 370 |
| Pop.[4] (residents) | | 1,760,000 | 550,000 | 170,000 |
| Density[4] (res./km$^2$) | | 4,500 | 1,230 | 490 |
| Vegetation[24] | % | 69.1 | 85.9 | 90.6 |
| | NDVI | 0.58 | 0.65 | 0.68 |
| Elevation* (m) | $\bar{x}$ | 30 | 117 | 232 |
| | Min | -14 | -3 | 125 |
| | Max | 229 | 629 | 476 |
| Latitude* | | 45.66318 | 46.98068 | 45.52405 |
| Longitude* | | -73.40981 | -71.13366 | -71.80254 |

*Topographic maps, 2024[1–3]

**Table S2.** Proportion of sequences (%) from the most relatively abundant (A) bacterial, (B) fungal, and (C) plant families across seasons and cities.

A) Bacteria (grey highlight showing four shared families identified as most relatively abundant taxa for all cities).

| **Family** | **All** | **Montréal** | | | | **Québec** | | | **Sherbrooke** | | | |
|---|---|---|---|---|---|---|---|---|---|---|---|---|
| | | SPR | SUM | FAL | TOT | SPR | FAL | TOT | SPR | SUM | FAL | TOT |
| *Nocardioidaceae* | 12.7 | 16.0 | 11.4 | 11.9 | 13.0 | 17.2 | 10.5 | 13.8 | 16.8 | 11.6 | 8.1 | 11.5 |
| *Geodermatophilaceae* | 9.8 | 8.0 | 9.7 | 11.3 | 9.9 | 9.7 | 13.8 | 11.8 | 7.1 | 6.7 | 9.5 | 8.0 |
| *Micrococacceae* | 7.6 | 14.1 | 14.3 | 6.6 | 10.6 | 6.9 | 4.2 | 5.5 | 6.3 | 5.8 | 4.0 | 5.2 |
| *Microbacteriaceae* | 5.9 | 6.6 | 4.6 | 5.0 | 5.4 | 6.0 | 5.0 | 5.5 | 6.0 | 4.7 | 9.0 | 6.8 |
| *Intrasporangiaceae* | 4.7 | 5.4 | 3.9 | 4.7 | 4.7 | 5.7 | 3.7 | 4.7 | 7.1 | 5.3 | 3.2 | 4.9 |
| *Hymenobacteriaceae* | 4.7 | 5.2 | 7.1 | 4.1 | 5.1 | 3.2 | 4.5 | 3.8 | 2.7 | 3.6 | 6.7 | 4.7 |
| *Deinococcaceae* | 3.8 | 2.2 | 7.4 | 6.0 | 5.2 | 2.2 | 2.7 | 2.5 | 1.2 | 2.4 | 4.2 | 2.9 |
| *Kineosporiaceae* | 3.2 | 1.6 | 3.0 | 4.3 | 3.2 | 1.2 | 4.7 | 2.9 | 1.3 | 1.5 | 6.0 | 3.3 |
| *Pseudonocardiaceae* | 2.4 | 1.8 | 1.6 | 3.2 | 2.4 | 1.8 | 2.2 | 2.0 | 1.9 | 4.5 | 1.9 | 2.7 |
| *Sphingomonadaceae* | 2.4 | 1.5 | 1.9 | 2.8 | 2.2 | 1.8 | 3.2 | 2.5 | 1.6 | 1.7 | 3.8 | 2.6 |
| *Beijerinckiaceae* | 2.3 | 0.7 | 0.9 | 2.5 | 1.6 | 1.0 | 4.4 | 2.7 | 0.9 | 1.9 | 5.2 | 3.0 |
| *Acetobacteraceae* | 1.9 | 1.5 | 1.3 | 1.7 | 1.5 | 2.4 | 2.1 | 2.3 | 2.1 | 1.8 | 2.2 | 2.0 |
| *Paracoccaceae* | 1.7 | 2.0 | 2.3 | 2.1 | 2.1 | 1.2 | 1.9 | 1.6 | 1.5 | 1.1 | 1.3 | 1.3 |
| *Oxalobacteraceae* | 1.7 | 1.8 | 2.4 | 1.4 | 1.8 | 1.8 | 1.5 | 1.6 | 1.4 | 1.3 | 1.7 | 1.5 |
| **Total (%)** | **64.7** | **68.4** | **71.8** | **67.5** | **68.8** | **62.1** | **64.5** | **63.3** | **57.8** | **54.0** | **66.8** | **60.3** |

B) Fungi (grey highlights showing three shared families identified as most relatively abundant taxa for all cities).

| Family | All | Montréal | | | | Québec | | | Sherbrooke | | | |
|---|---|---|---|---|---|---|---|---|---|---|---|---|
| | | SPR | SUM | FAL | TOT | SPR | FAL | TOT | SPR | SUM | FAL | TOT |
| *Cladosporiaceae* | 21.6 | 19.6 | 32.8 | 38.5 | 31.6 | 17.7 | 12.5 | 15.1 | 6.6 | 11.7 | 16.5 | 12.1 |
| *Polyporaceae* | 13.1 | 9.9 | 3.9 | 14.9 | 10.8 | 17.7 | 18.6 | 18.1 | 12.5 | 4.1 | 19.7 | 12.1 |
| *Fomitopsidaceae* | 12.2 | 18.6 | 0.1 | 2.3 | 6.6 | 9.5 | 21.1 | 15.3 | 38.3 | 1.7 | 19.6 | 18.2 |
| *Phanerochaetaceae* | 2.8 | 5.4 | 1.9 | 1.0 | 2.5 | 5.1 | 0.4 | 2.7 | 6.3 | 3.7 | 0.5 | 3.2 |
| *Pleosporaceae* | 2.7 | 3.0 | 9.0 | 3.5 | 4.7 | 2.2 | 1.4 | 1.8 | 0.5 | 0.4 | 1.2 | 0.7 |
| *Irpicaceae* | 2.5 | 2.5 | 7.8 | 2.6 | 3.8 | 0.0 | 0.3 | 0.2 | 0.0 | 6.5 | 0.6 | 2.6 |
| *Cortinariaceae* | 2.4 | 0.0 | 0.0 | 1.8 | 0.8 | 0.0 | 10.7 | 5.4 | 0.0 | 0.0 | 5.4 | 2.0 |
| *Cerrenaceae* | 2.2 | 5.5 | 1.7 | 0.1 | 2.1 | 2.1 | 0.1 | 1.1 | 1.3 | 8.3 | 0.1 | 3.4 |
| *Mycosphaerellaceae* | 2.2 | 0.1 | 0.8 | 0.3 | 0.3 | 9.8 | 0.0 | 4.9 | 0.2 | 6.8 | 0.1 | 2.6 |
| *Discinellaceae* | 2.1 | 1.6 | 4.9 | 0.0 | 1.6 | 2.1 | 0.0 | 1.1 | 0.8 | 10.1 | 0.0 | 3.9 |
| *Peniophoraceae* | 1.9 | 0.7 | 7.1 | 3.0 | 3.3 | 0.2 | 0.3 | 0.3 | 0.0 | 0.6 | 3.1 | 1.4 |
| *Hydnodontaceae* | 1.6 | 1.3 | 0.7 | 2.2 | 1.6 | 1.4 | 1.1 | 1.3 | 2.7 | 1.6 | 1.8 | 2.0 |
| *Didymellaceae* | 1.6 | 2.3 | 1.4 | 1.7 | 1.8 | 1.5 | 1.2 | 1.4 | 0.9 | 1.5 | 1.9 | 1.5 |
| *Bulleribasidiaceae* | 1.5 | 3.8 | 0.3 | 1.3 | 1.8 | 2.6 | 1.9 | 2.2 | 0.6 | 0.1 | 0.5 | 0.4 |
| **Total (%)** | **70.5** | **74.2** | **72.4** | **73.2** | **73.3** | **72.1** | **69.4** | **70.8** | **70.8** | **57.2** | **71.1** | **66.0** |

C) Plant (grey highlights showing the shared family identified as most relatively abundant taxa for all cities).

| Family | All | Montréal | | | | Québec | | | Sherbrooke | | | |
|---|---|---|---|---|---|---|---|---|---|---|---|---|
| | | SPR | SUM | FAL | TOT | SPR | FAL | TOT | SPR | SUM | FAL | TOT |
| *Pinaceae* | 35.0 | 0.2 | 98.3 | 21.5 | 33.3 | 5.0 | 41.7 | 23.4 | 0.4 | 88.3 | 42.0 | 47.0 |
| *Asteraceae* | 13.9 | 0.1 | 0.0 | 59.6 | 28.0 | 0.3 | 5.9 | 3.1 | 0.1 | 0.6 | 6.3 | 2.9 |
| *Oleaceae* | 9.8 | 26.0 | 0.2 | 0.9 | 8.2 | 34.7 | 1.7 | 18.2 | 10.1 | 2.6 | 4.1 | 5.1 |
| *Sapindaceae* | 9.8 | 52.8 | 0.1 | 0.6 | 15.8 | 5.2 | 2.4 | 3.8 | 21.0 | 0.9 | 1.3 | 6.1 |
| *Betulaceae* | 8.7 | 7.5 | 0.1 | 0.5 | 2.5 | 21.1 | 2.7 | 11.9 | 50.4 | 0.1 | 5.4 | 14.9 |
| *Poaceae* | 3.0 | 0.1 | 0.1 | 1.9 | 0.9 | 0.0 | 9.5 | 4.8 | 0.1 | 1.8 | 9.3 | 4.5 |
| *Equisetaceae* | 2.5 | 7.2 | 0.1 | 0.0 | 2.1 | 5.5 | 0.2 | 2.8 | 10.3 | 0.0 | 0.0 | 2.6 |
| **Total (%)** | **82.6** | **93.9** | **98.8** | **85.1** | **90.9** | **71.8** | **64.1** | **68.0** | **92.4** | **94.3** | **68.5** | **83.1** |

**Table S3.** Core microbiome ASVs taxonomy, prevalence and mean relative abundance. Each row represents a distinct ASVs. Core ASVs are present in at least 95 % of samples with a mean relative abundance of at least 1 % across the samples in which they are present.

| **Taxonomy** | | | | | **Relative abundance** | | **Preva-lence** |
|---|---|---|---|---|---|---|---|
| **Class** | **Order** | **Family** | **Genus** | **Species** | **Mean** | **SD** | **(%)** |
| **BACTERIA** | | | | | | | |
| ***Actinobacteria*** | *Propionibacteriales* | *Nocardioidaceae* | *Nocardioides* | *aquaticus* | 0.015 | 0.012 | 100 |
| ***Actinobacteria*** | *Propionibacteriales* | *Nocardioidaceae* | *Nocardioides* | unclassified | 0.021 | 0.009 | 100 |
| ***Actinobacteria*** | *Frankiales* | *Unclassified* | *Unclassified* | unclassified | 0.020 | 0.011 | 100 |
| ***Actinobacteria*** | *Frankiales* | *Geodermatophilaceae* | *Blastococcus* | *aggregatus* | 0.019 | 0.015 | 99.3 |
| ***Actinobacteria*** | *Frankiales* | *Geodermatophilaceae* | *Blastococcus* | unclassified | 0.021 | 0.016 | 100 |
| ***Actinobacteria*** | *Frankiales* | *Geodermatophilaceae* | *Blastococcus* | unclassified | 0.019 | 0.016 | 100 |
| ***Actinobacteria*** | *Micrococcales* | *Micrococcaceae* | *Arthrobacter* | unclassified | 0.023 | 0.025 | 99.3 |
| ***Actinobacteria*** | *Micrococcales* | *Micrococcaceae* | *Kocuria* | unclassified | 0.015 | 0.022 | 95.4 |
| **FUNGI** | | | | | | | |
| ***Dothideomycetes*** | *Cladosporiales* | *Cladosporiaceae* | *Cladosporium* | *herbarum* | 0.077 | 0.102 | 100 |
| ***Dothideomycetes*** | *Cladosporiales* | *Cladosporiaceae* | *Cladosporium* | unclassified | 0.096 | 0.140 | 99.3 |
| ***Dothideomycetes*** | *Pleosporales* | *Pleosporaceae* | *Alternaria* | *alternata* | 0.020 | 0.032 | 97.3 |
| ***Agaricomycetes*** | *Trechisporales* | *Hydnodontaceae* | *Sistotremastrum* | *guttuliferum* | 0.012 | 0.014 | 100 |
| ***Agaricomycetes*** | *Polyporales* | *Polyporaceae* | *Trametes* | *versicolor* | 0.025 | 0.021 | 100 |
| ***Agaricomycetes*** | *Hymenochaetales* | *Rickenellaceae* | *Resinicium* | *bicolor* | 0.017 | 0.028 | 96.6 |
| ***Agaricomycetes*** | *Polyporales* | *Phanerochaetaceae* | *Bjerkandera* | *adusta* | 0.013 | 0.016 | 95.3 |
| ***Agaricomycetes*** | *Polyporales* | *Phanerochaetaceae* | *Bjerkandera* | unclassified | 0.012 | 0.014 | 95.9 |
| **POLLEN** | | | | | | | |
| ***Magnoliopsida*** | *Lamiales* | *Oleaceae* | *Syringa* | unclassified | 0.112 | 0.211 | 100 |

**Table S4.** Mean temperatures and precipitations for sampling periods per cities in 2022.

| Climatic variables | | MTL | QUB | SHB |
|---|---|---|---|---|
| Spring | Mean temperature (°C) | 9.4 | 6.6 | 14.0 |
| | Mean precipitations (mm) | 1.5 | 2.0 | 3.4 |
| Summer | Mean temperature (°C) | 18.1 | - | 15.1 |
| | Mean precipitations (mm) | 5.0 | - | 4.8 |
| Fall | Mean temperature (°C) | 15.2 | 14.0 | 12.8 |
| | Mean precipitations (mm) | 6.5 | 2.1 | 5.2 |
| Global | Max temperature (°C) | 31.2 | 28.0 | 31.4 |
| | Min temperature (°C) | 1.2 | 0.4 | -2.4 |

Public data from Environment and Climate Change Canada[5]

**Table S5.** Primers used for qPCRs and sequencing.

| Marker | Technique | Primer | Sequence | Reference |
|---|---|---|---|---|
| 16S V5-V6 | qPCR & Sequencing | 799F | 5'–AACMGGATTAGATACCCKG-3' | Aydogan et al. 2018 |
| | Sequencing | 1115R | 5'–AGGGTTGCGCTCGTTG–3' | |
| 16S | qPCR | 1193R | 5'-ACGTCATCCCCACCTTCC-3' | Bodenhausen et al., 2013 |
| ITS | qPCR & Sequencing | ITS1F | 5'–CTTGGTCATTTAGAGGAAGTAA–3' | Gardes & Bruns 1993 |
| | | ITS2 | 5'–GCTGCGTTCTTCATCGATGC–3' | White et al. 1990 |
| *trnL* | qPCR & Sequencing | c-A49325 | 5'–CGAAATCGGTAGACGCTACG–3' | Taberlet et al. 2007 |
| | | d-B49863 | 5'–GGGGATAGAGGGACTTGAAC–3' | |

**Table S6.** Summary statistics of sample ASVs across each marker*.

| Marker | n | Reads | ASVs | Sequences per sample $\bar{x}$ (range) | SD | ASVs per sample $\bar{x}$ (range) | SD | ASV prevalence $\bar{x}$ (range) | SD |
|---|---|---|---|---|---|---|---|---|---|
| Bacteria | 152 | 3,527,682 | 5,896 | 23,208 (9,358 – 41,745) | 5,744 | 1,181 (300 – 2,170) | 415 | 30 (1 – 152) | 31 |
| Fungi | 148 | 1,377,622 | 1,348 | 9,308 (2,625 – 48,749) | 5,193 | 267 (121 – 427) | 63 | 29 (1 – 148) | 25 |
| Pollen | 147 | 1,665,375 | 378 | 11,329 (2,543 – 30,358) | 6,441 | 54 (15 – 93) | 18 | 21 (1 – 147) | 22 |

*After filtering for quality, chimeras, and removing ASVs with fewer than 50 sequences. ASV prevalence is the number of samples in which an ASV is found.

**Table S7.** Differential abundance analysis. LFC: Natural log (ln) fold-change. Showing only genera with a $p_{adj.}$ <0.01 having passed the pseudocount sensitivity test (a total of 586 genera were tested twice) with a LFC>1.5. Only LFC with p<0.01 are reported. *Season* represents which season the genus is enriched (or depleted, if LFC<0) compared to summer. P-values were adjusted using the Holm procedure. **Genus Incertae sedis*

| Dataset | Genus | Season | LFC | Standard error | Adj. p-value |
|---|---|---|---|---|---|
| Bacteria | Aureimonas | Fall | 2.687 | 0.303 | 3.8e-12 |
| Bacteria | Cryobacterium | Fall | -1.548 | 0.287 | 3.1e-04 |
| Bacteria | Exiguobacterium | Spring | -2.592 | 0.334 | 6.4e-09 |
| Bacteria | Frigoribacterium | Fall | 1.774 | 0.307 | 5.4e-05 |
| Bacteria | Methylobacterium | Fall | 1.408 | 0.247 | 7.2e-05 |
| Bacteria | Paenarthrobacter | Fall | -1.393 | 0.288 | 4.0e-03 |
| Bacteria | Pseudokineococcus | Fall | 1.806 | 0.303 | 3.0e-05 |
| Fungi | Amphinema | Fall | 5.090 | 0.352 | 4.9e-16 |
| Fungi | Aspergillus | Spring | 1.677 | 0.336 | 2.3e-03 |
| Fungi | Bullera | Fall | 2.365 | 0.324 | 7.6e-08 |
| Fungi | Ceratobasidiaceae* | Spring | -1.922 | 0.383 | 9.7e-03 |
| Fungi | Ceratobasidiaceae* | Fall | -2.508 | 0.384 | 5.4e-05 |

| Dataset | Genus | Season | LFC | Standard error | Adj. p-value |
|---|---|---|---|---|---|
| Fungi | Ceriporia | Fall | 1.925 | 0.337 | 5.8e-03 |
| Fungi | Cerrena | Fall | -2.564 | 0.428 | 2.7e-05 |
| Fungi | Coprinopsis | Spring | -2.693 | 0.353 | 1.0e-05 |
| Fungi | Crustomyces | Spring | -1.812 | 0.344 | 4.7e-03 |
| Fungi | Daedaleopsis | Spring | 2.346 | 0.350 | 4.0e-06 |
| Fungi | Deconica | Spring | -1.867 | 0.333 | 5.5e-03 |
| Fungi | Exidia | Spring | -2.833 | 0.410 | 6.3e-07 |
| Fungi | Exidia | Fall | -3.604 | 0.392 | 9.7e-12 |
| Fungi | Fomes | Fall | -2.383 | 0.392 | 1.9e-05 |
| Fungi | Fomitopsis | Spring | 3.597 | 0.403 | 2.5e-12 |
| Fungi | Fomitopsis | Fall | 2.976 | 0.421 | 8.0e-08 |
| Fungi | Fraxinicola | Fall | -2.574 | 0.442 | 2.1e-03 |
| Fungi | Fuscoporia | Spring | -1.917 | 0.314 | 3.3e-05 |
| Fungi | Ganoderma | Spring | -2.436 | 0.320 | 3.0e-08 |
| Fungi | Ganoderma | Fall | 2.851 | 0.337 | 5.5e-10 |
| Fungi | Gloeocystidiellum | Spring | -1.940 | 0.343 | 1.1e-03 |
| Fungi | Gloeophyllum | Fall | -2.857 | 0.371 | 4.0e-09 |
| Fungi | Gloeoporus | Spring | -1.589 | 0.306 | 1.4e-03 |
| Fungi | Gloeoporus | Fall | 1.788 | 0.334 | 7.2e-04 |
| Fungi | Gymnopus | Spring | -3.454 | 0.389 | 7.6e-09 |
| Fungi | Gymnopus | Fall | -2.330 | 0.402 | 4.9e-04 |
| Fungi | Hannaella | Fall | 2.874 | 0.346 | 9.5e-09 |
| Fungi | Hyphoderma | Spring | -2.000 | 0.310 | 1.7e-05 |

| Dataset | Genus | Season | LFC | Standard error | Adj. p-value |
|---|---|---|---|---|---|
| Fungi | Hyphodermella | Spring | -2.445 | 0.378 | 1.7e-04 |
| Fungi | Hyphodontia | Fall | 1.952 | 0.378 | 1.1e-03 |
| Fungi | Hypoxylon | Spring | 1.943 | 0.322 | 6.6e-04 |
| Fungi | Irpex | Spring | -3.715 | 0.488 | 1.5e-07 |
| Fungi | Irpex | Fall | -3.389 | 0.467 | 6.6e-07 |
| Fungi | Lachnum | Spring | -1.869 | 0.358 | 3.3e-03 |
| Fungi | Lachnum | Fall | -2.908 | 0.355 | 5.5e-08 |
| Fungi | Leptospora | Spring | -2.753 | 0.396 | 1.3e-05 |
| Fungi | Leptospora | Fall | -2.889 | 0.395 | 3.5e-06 |
| Fungi | Mollisia | Spring | -2.204 | 0.319 | 2.5e-06 |
| Fungi | Mollisia | Fall | -2.234 | 0.325 | 3.0e-06 |
| Fungi | Mrakia | Spring | 1.876 | 0.324 | 2.4e-04 |
| Fungi | Osteina | Fall | 1.834 | 0.312 | 1.7e-04 |
| Fungi | Oxyporus | Spring | -1.765 | 0.364 | 3.8e-03 |
| Fungi | Perenniporia | Fall | 2.916 | 0.319 | 4.8e-12 |
| Fungi | Postia | Fall | 2.646 | 0.321 | 3.0e-08 |
| Fungi | Punctularia | Fall | -2.021 | 0.331 | 2.2e-03 |
| Fungi | Ramularia | Spring | -2.350 | 0.435 | 4.9e-04 |
| Fungi | Ramularia | Fall | -3.536 | 0.435 | 1.2e-09 |
| Fungi | Steccherinum | Spring | -2.842 | 0.404 | 1.5e-06 |
| Fungi | Teratosphaeria | Spring | -2.742 | 0.447 | 1.5e-03 |
| Fungi | Teratosphaeria | Fall | -2.780 | 0.453 | 1.4e-03 |
| Fungi | Trametes | Fall | 2.203 | 0.350 | 4.3e-06 |

| Dataset | Genus | Season | LFC | Standard error | Adj. p-value |
|---|---|---|---|---|---|
| Fungi | Trichaptum | Spring | -2.283 | 0.457 | 4.3e-03 |
| Fungi | Tubulicrinis | Spring | -2.156 | 0.382 | 2.3e-04 |
| Fungi | Venturia | Fall | -2.024 | 0.346 | 3.4e-04 |
| Fungi | Vishniacozyma | Spring | 1.918 | 0.355 | 3.4e-04 |
| Fungi | Xenasmatella | Fall | 2.486 | 0.348 | 8.8e-08 |
| Pollen | Acer | Spring | 4.489 | 0.403 | 9.8e-18 |
| Pollen | Betula | Spring | 4.368 | 0.424 | 9.1e-16 |
| Pollen | Dipteronia | Spring | 3.109 | 0.366 | 8.4e-09 |
| Pollen | Equisetum | Spring | 4.494 | 0.405 | 4.6e-15 |
| Pollen | Fraxinus | Spring | 2.538 | 0.349 | 1.5e-04 |
| Pollen | Grimmia | Spring | 2.228 | 0.362 | 1.6e-03 |
| Pollen | Juniperus | Spring | 3.097 | 0.459 | 2.4e-06 |
| Pollen | Pinus | Spring | -8.350 | 0.458 | 4.6e-35 |
| Pollen | Pinus | Fall | -4.632 | 0.528 | 8.3e-12 |
| Pollen | Syringa | Spring | 3.952 | 0.432 | 6.5e-13 |

**Figure S1.** Distribution of sites along vegetation cover and median family income gradients for each site per city. The vegetation cover gradient (NDVI) is categorized in three bins (1: 0–0.3, 2: 0.3–0.5, 3: 0.5–1) and the median household income gradient is categorized in four bins (1: < 40k CA$, 2: 40k–70k CA$, 3: 70k–100k CA$, 4: > 100k CA$).

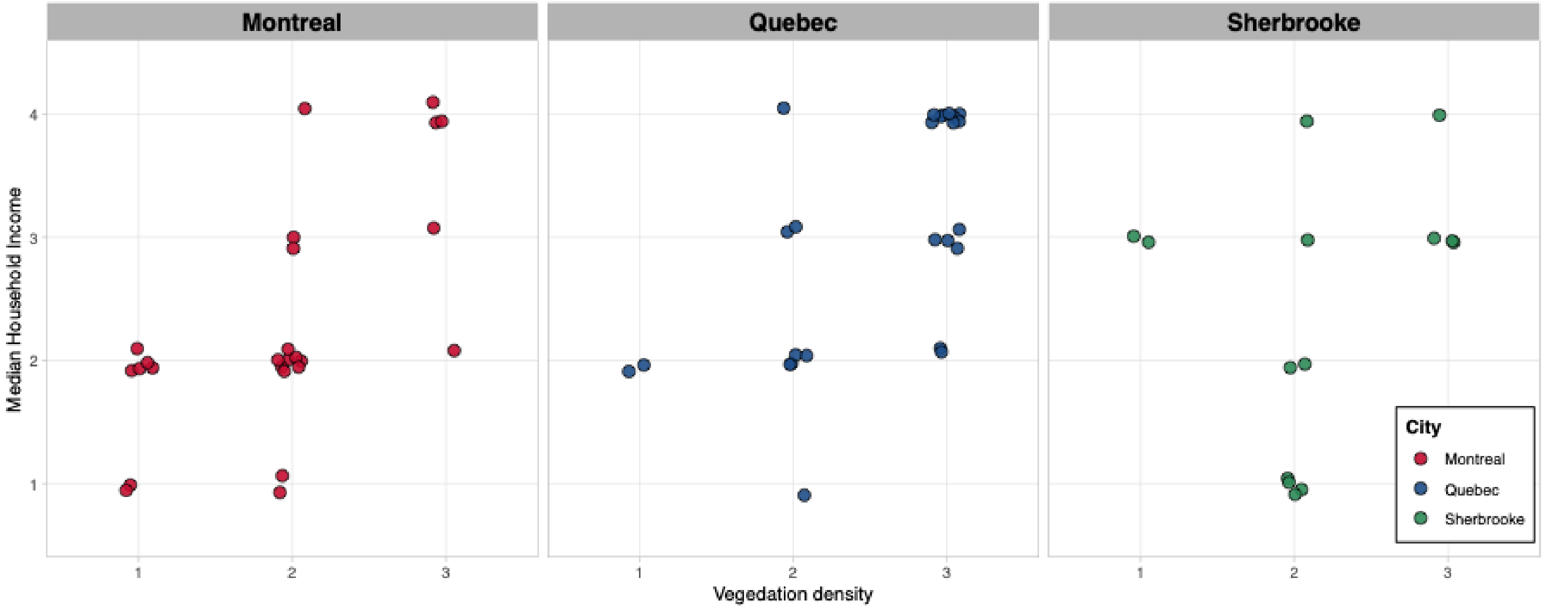

**Figure S2.** Temperature recordings per sampling dates per city from May to October 2022 showing (A) Montréal, (B) Québec City, and (C) Sherbrooke.

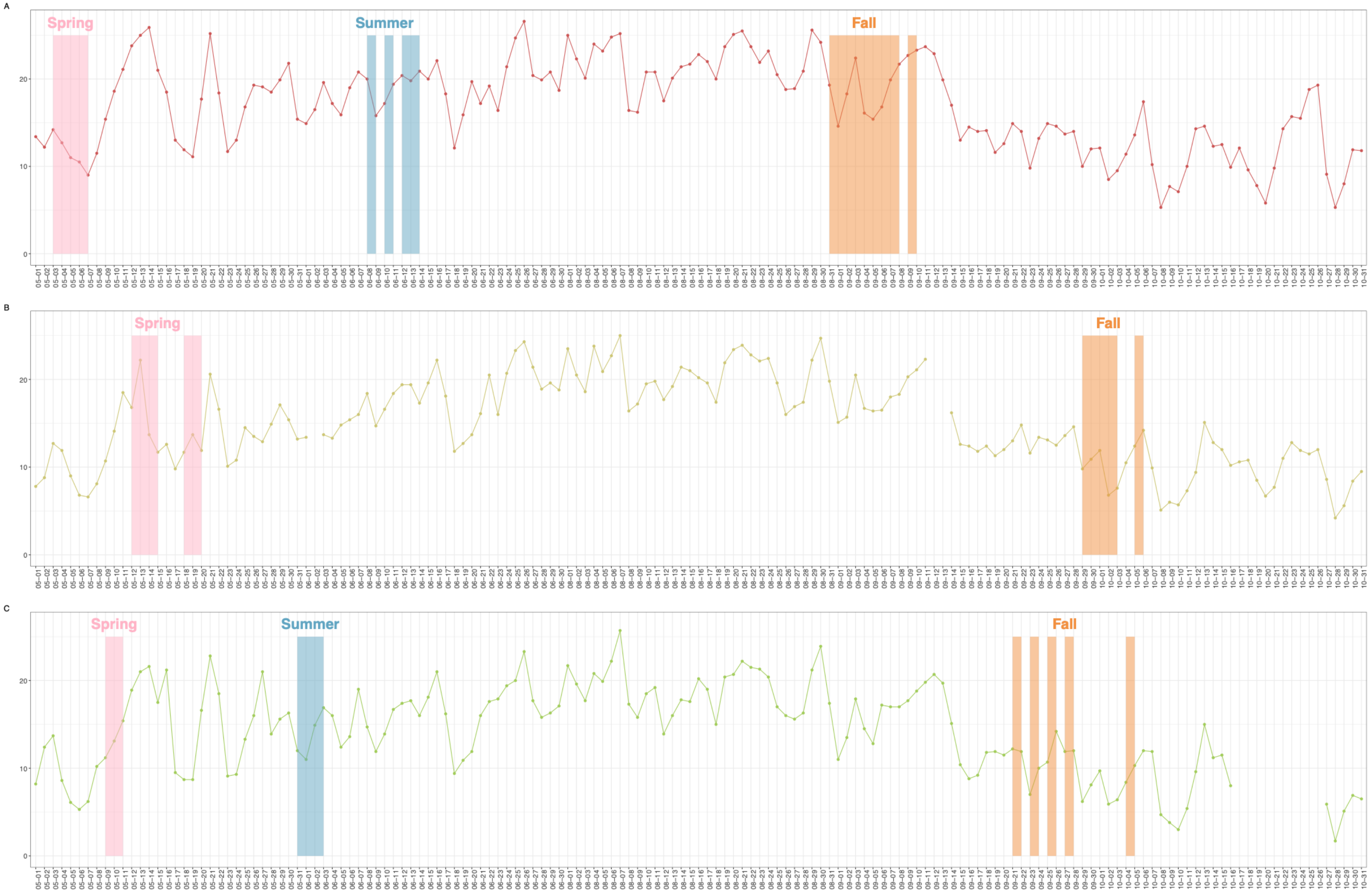

**Figure S3.** Precipitations recordings per sampling dates per city from May to October 2022 showing (A) Montréal, (B) Québec City, and (C) Sherbrooke.

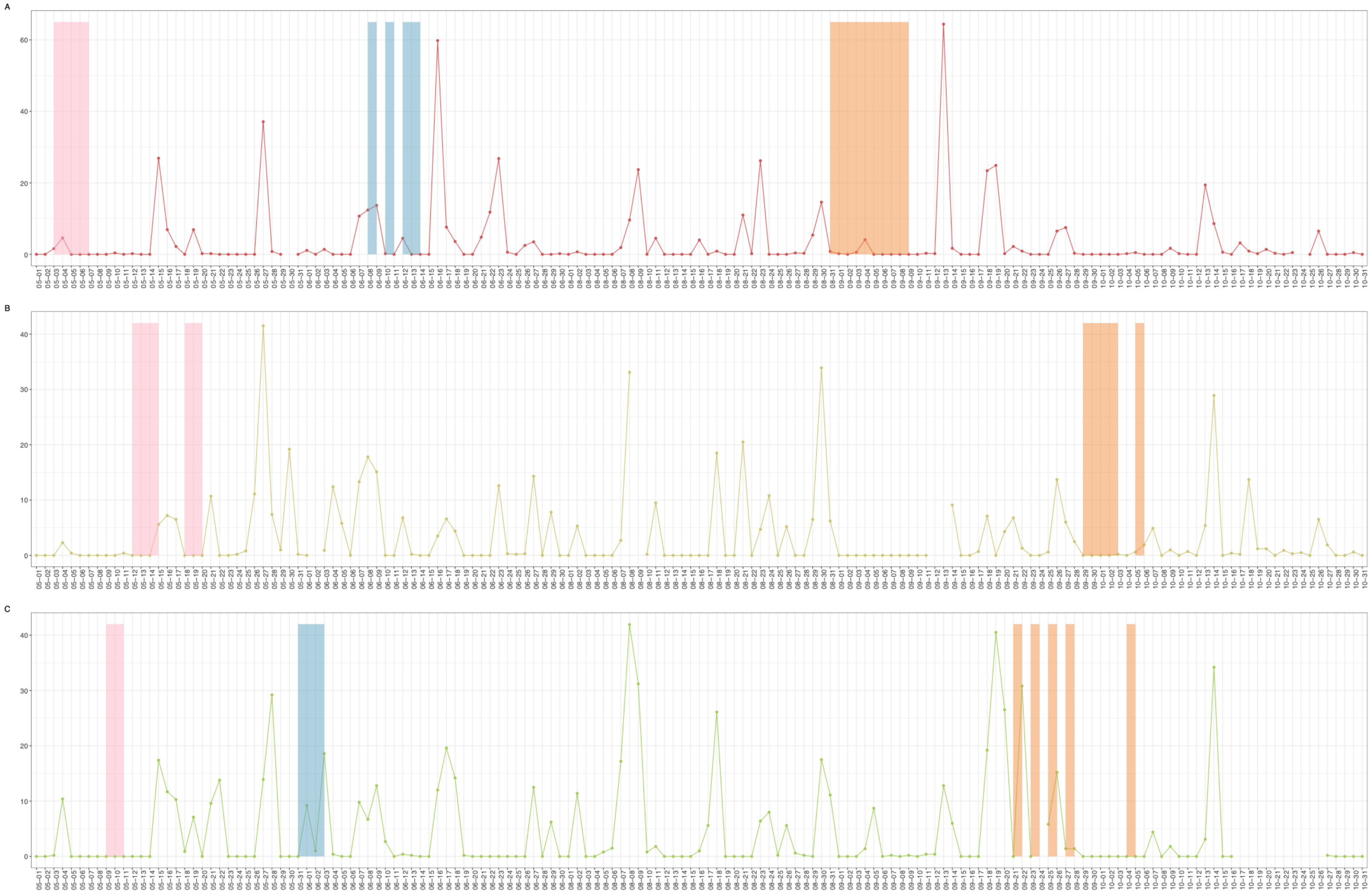

**Figure S4.** Percent contribution of turnover and nestedness to changes in bioaerosol community structure across three taxa, bacteria, fungi, and pollen, in urban environments. Panels A–C show results for Montréal, D–F for Québec City, and G–I for Sherbrooke. Each panel illustrates the relative influence of species turnover (community replacement) vs. nestedness (species loss or gain without replacement) in shaping bioaerosol diversity across sampling sites within each city.

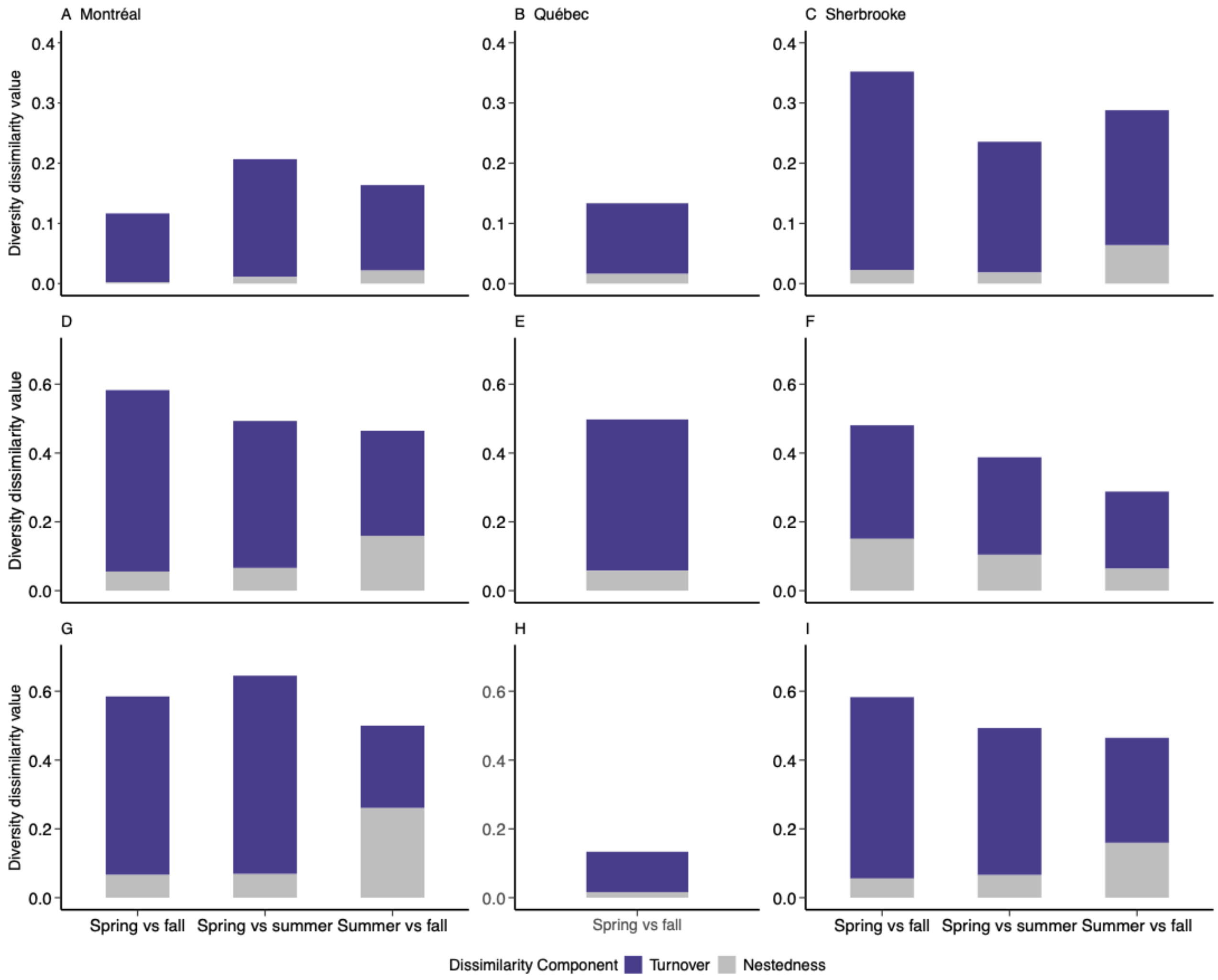

**Figure S5.** Relative abundance of bacterial families per sample across sampling periods and cities.

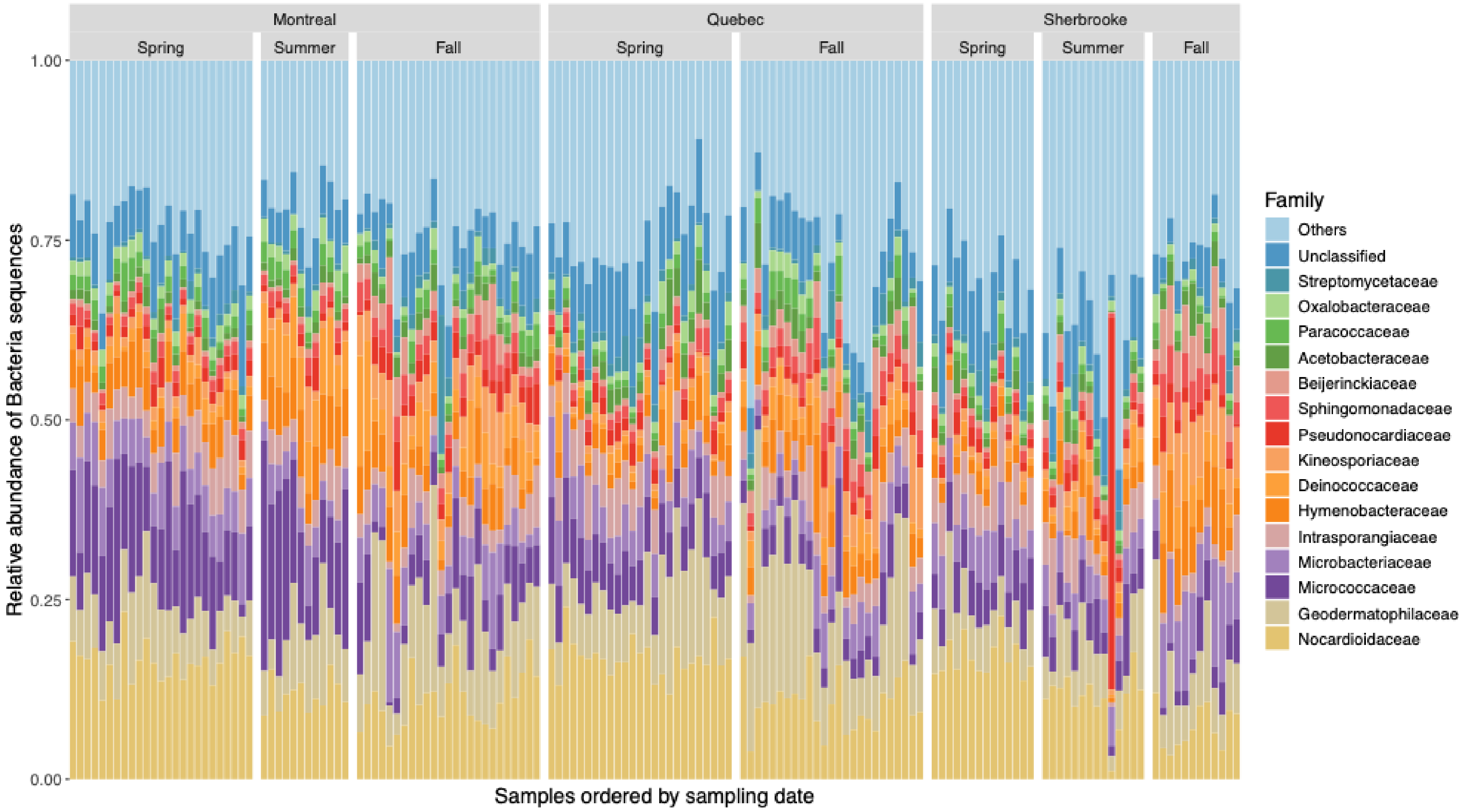

51 **Figure S6.** Relative abundance of fungal families per sample across sampling periods and cities.

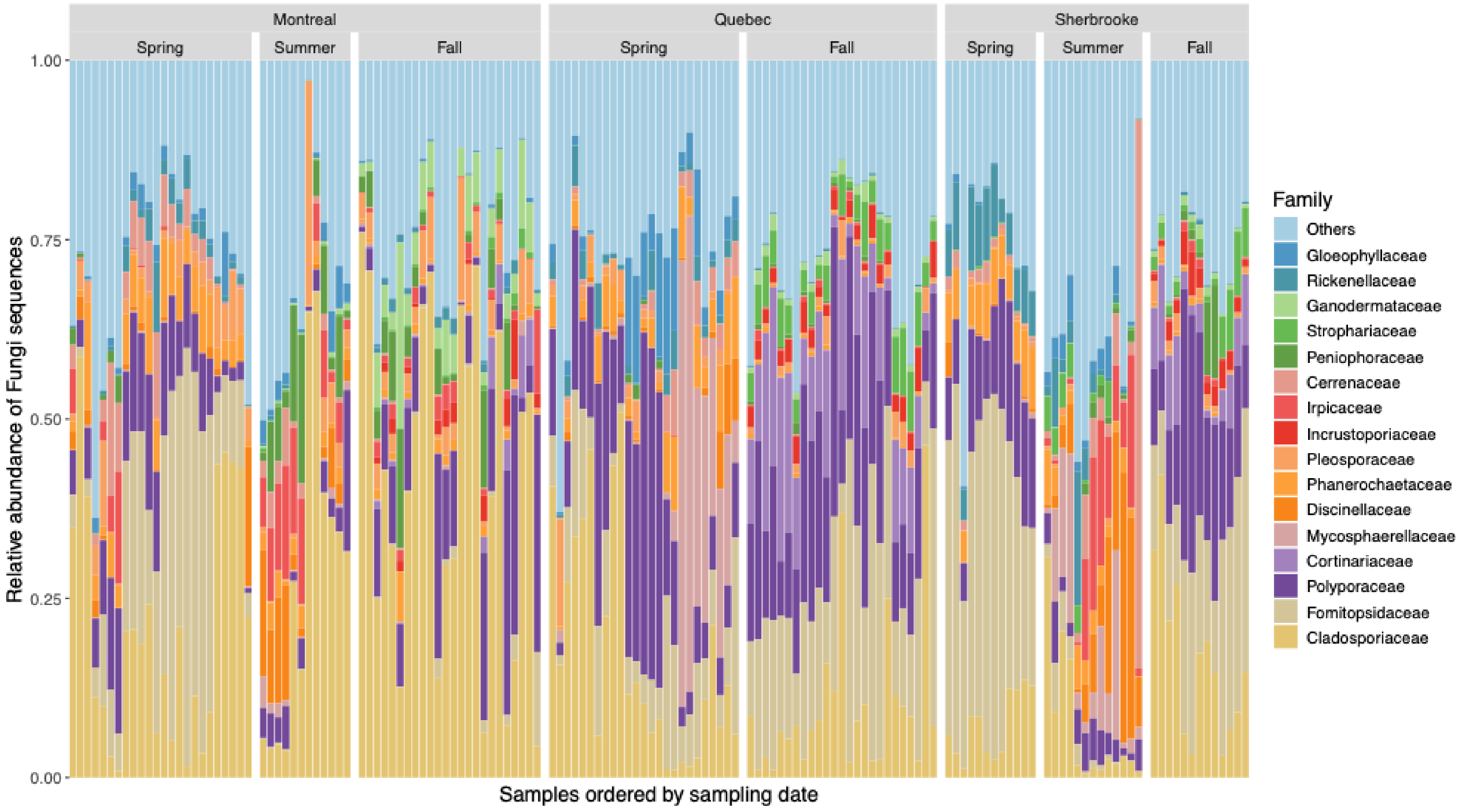


52
53

**Figure S7.** Relative abundance of plant (pollen) families per sample across sampling periods and cities.

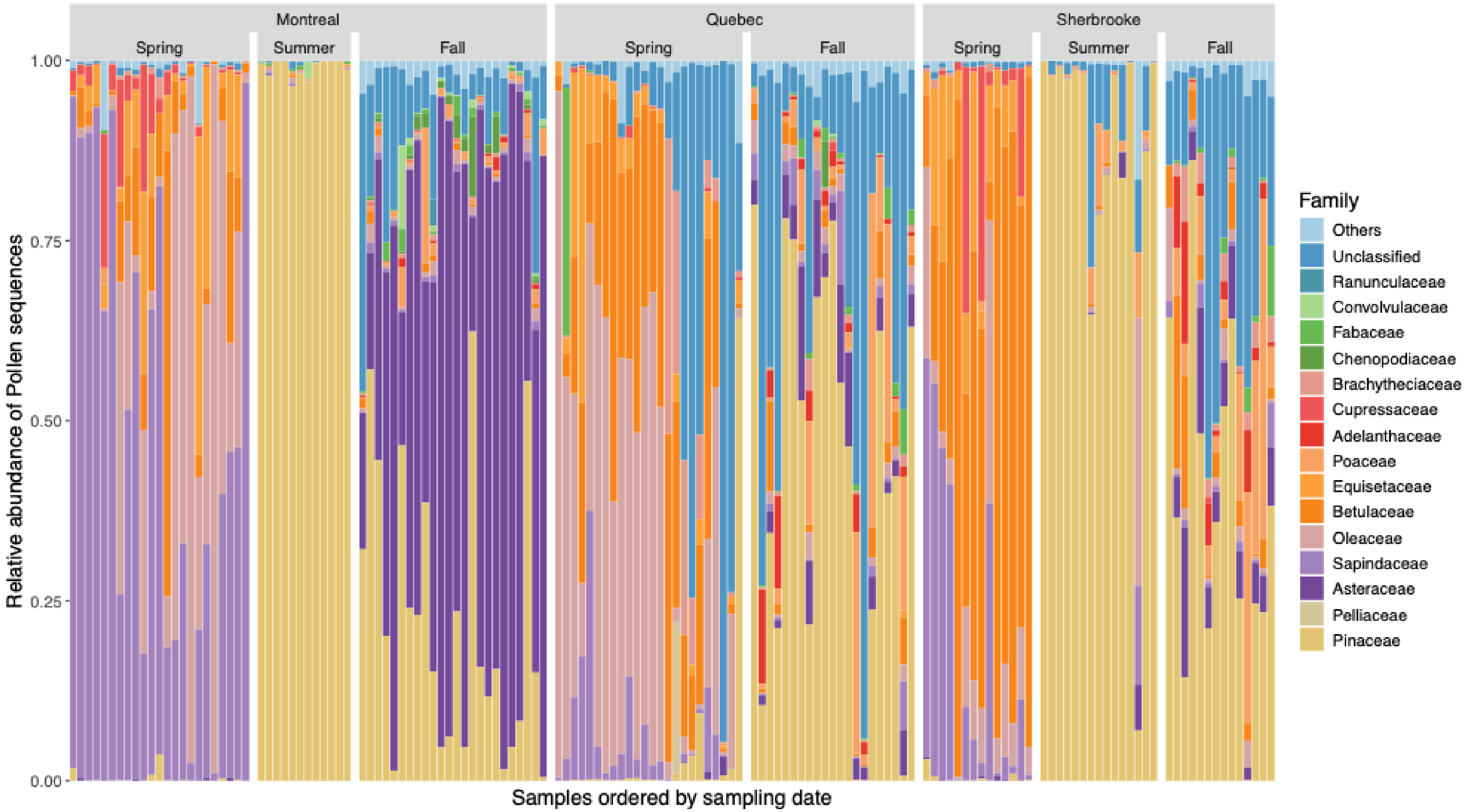

Figure S8. Distribution of Bray-Curtis dissimilarities between samples at given time and cities for each amplicon.

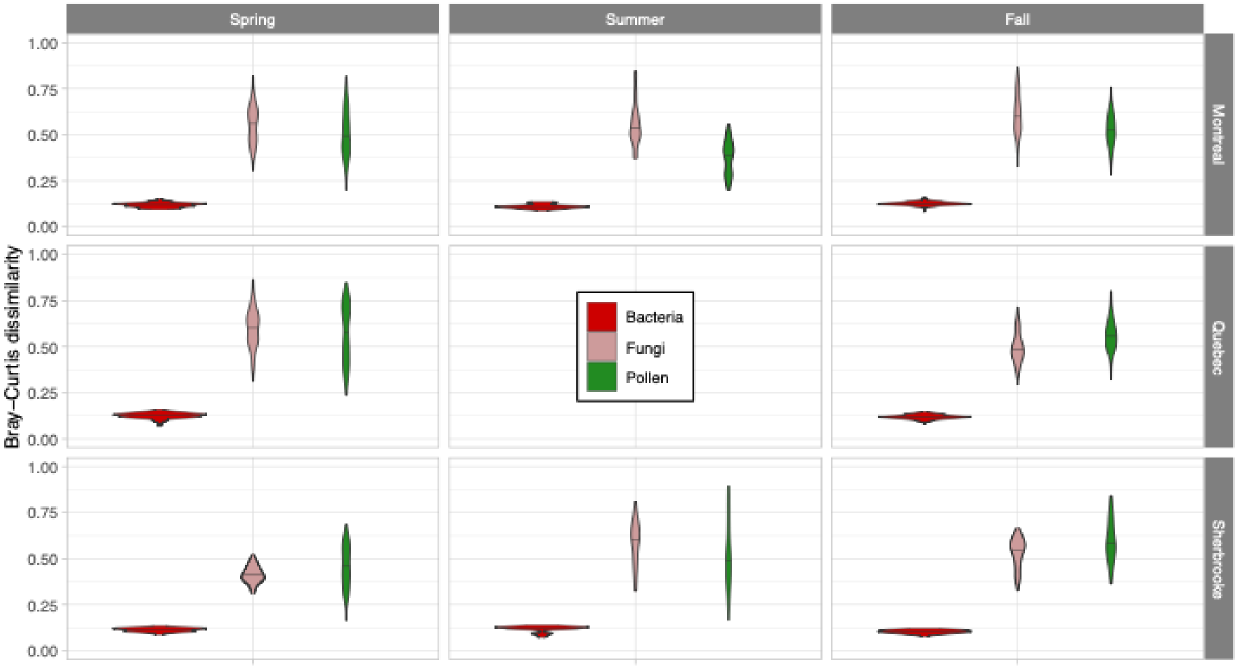

**Figure S9.** Alpha diversity across (A) sampling periods, (B) vegetation index (NDVI), and (C) median household income for bacterial, fungal, and plant bioaerosols for all samples from all three cities. Statistical significance was assessed using the Wilcoxon signed-rank test with p-value correction using the Holm procedure; asterisks indicate significant differences (*p < 0.05, **p < 0.01, ***p < 0.001, ****p < 0.0001).

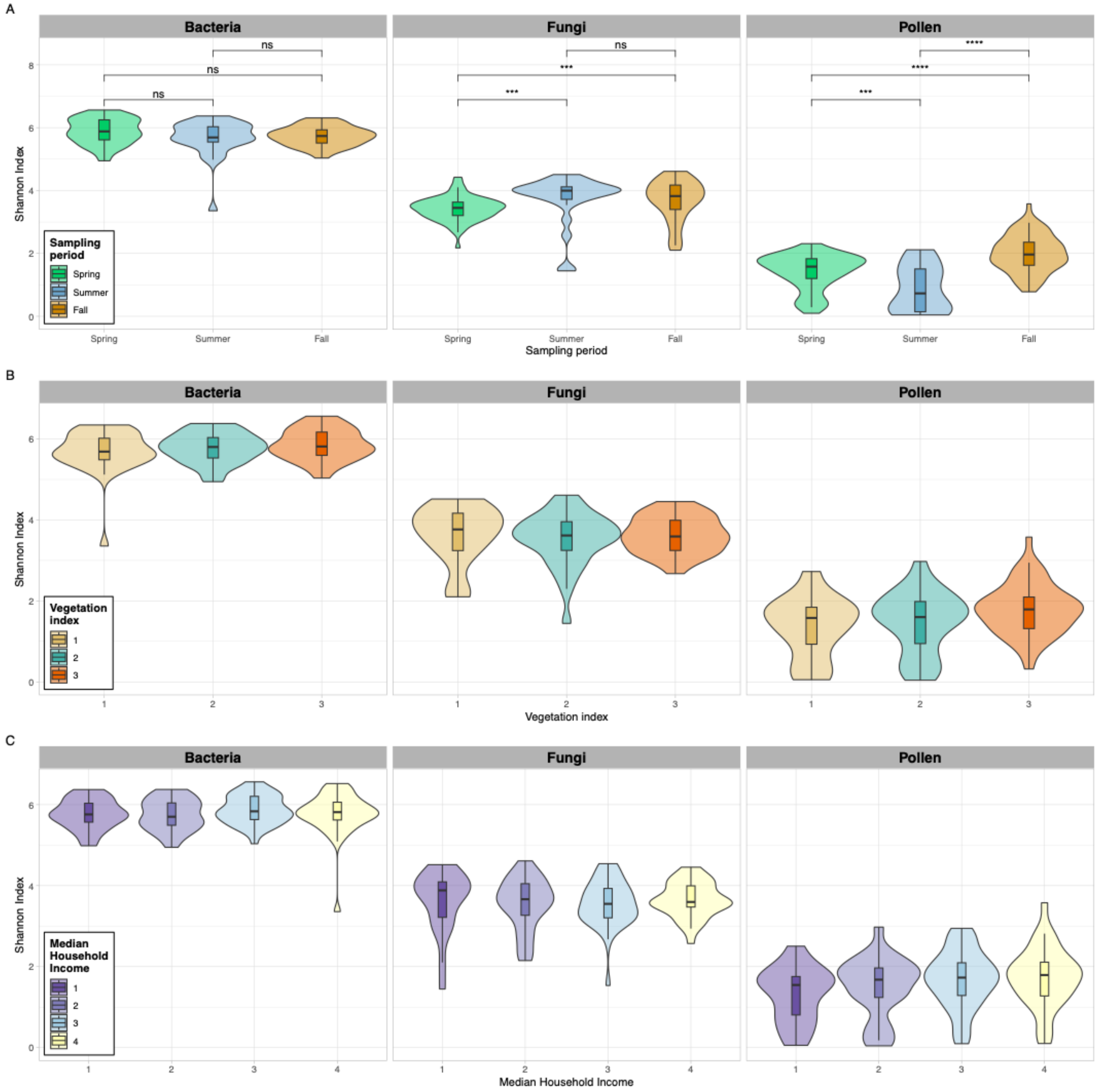

**Figure S10.** Alpha diversity across sampling periods (A) spring, (B) summer, and (C) fall for bacterial, fungal, and plant bioaerosols for all samples across the three cities. Statistical significance was assessed using the Wilcoxon signed-rank test with p-value correction using the Holm procedure; asterisks indicate significant differences (*p < 0.05, **p < 0.01, ***p < 0.001, ****p < 0.0001).

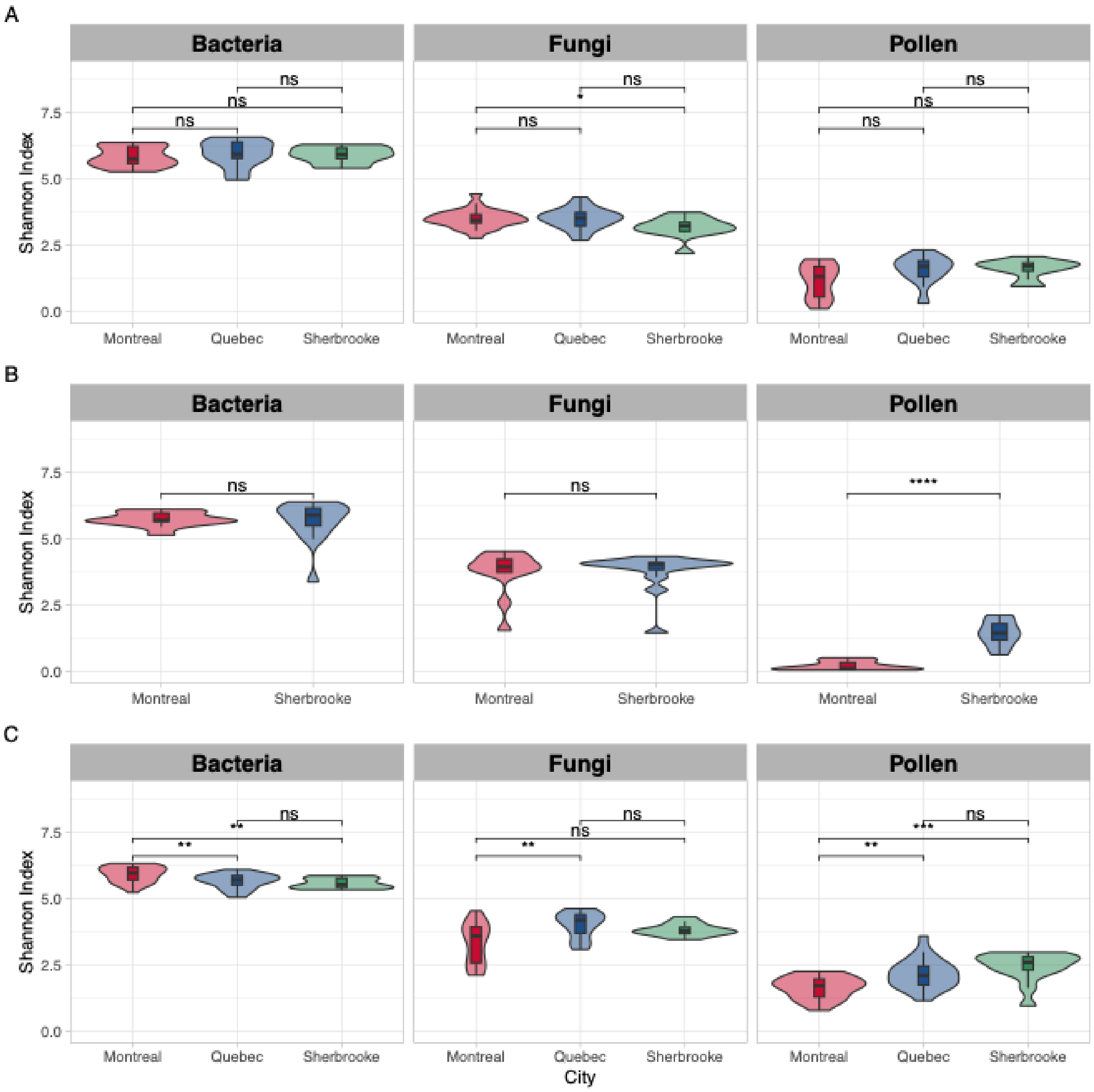

**Figure S11.** Alpha diversity across vegetation cover (NDVI) and median household income in Montréal. The vegetation cover gradient (NDVI) is categorized in three bins (1: 0–0.3, 2: 0.3–0.5, 3: 0.5–1) and the median household income gradient is categorized in four bins (1: < 40k CA$, 2: 40k–70k CA$, 3: 70k–100k CA$, 4: > 100k CA$). There was no statistical significance detected using the Wilcoxon signed-rank test with p-value correction using the Holm procedure.

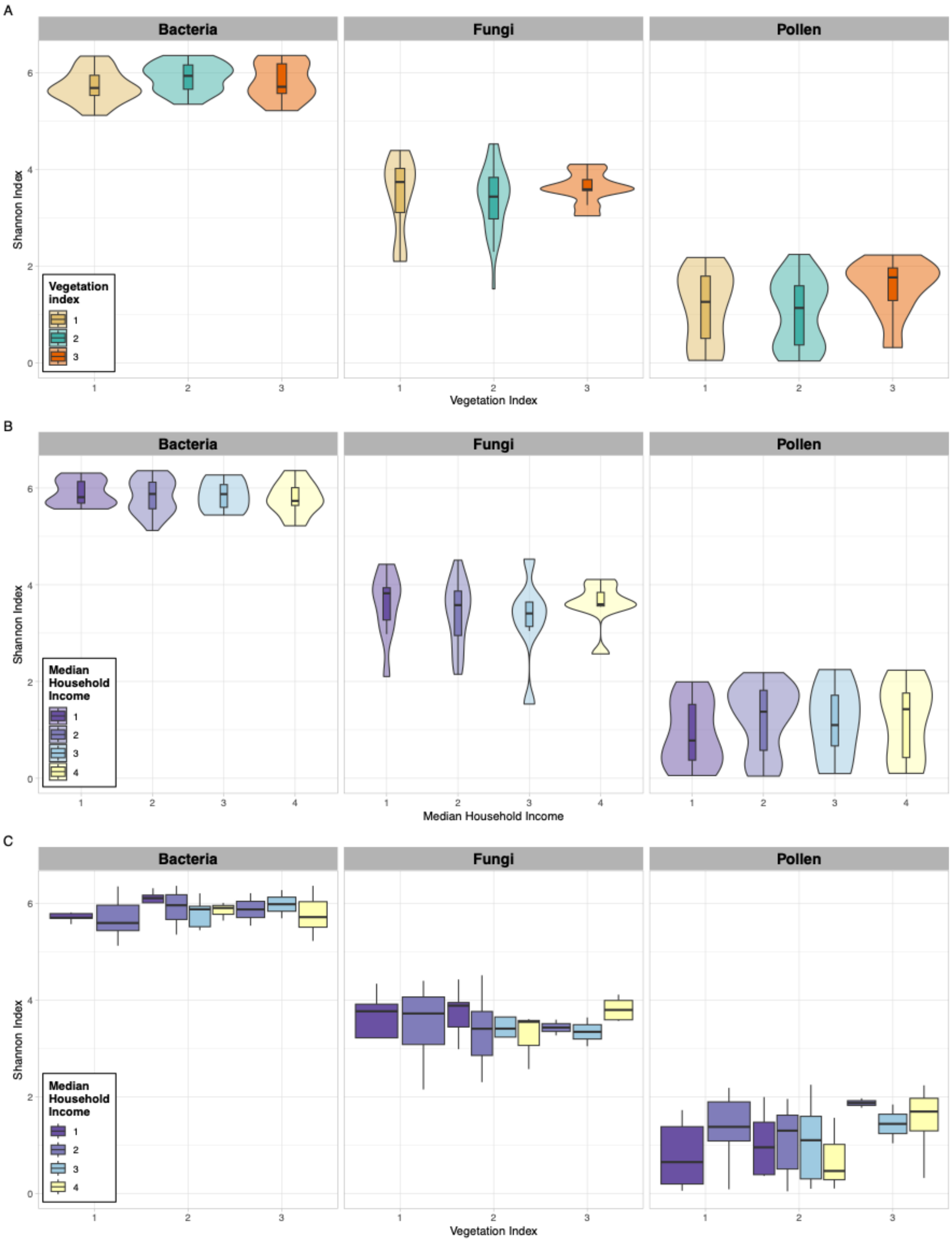

**Figure S12.** Alpha diversity across vegetation cover (NDVI) and median household income in Québec. The vegetation cover gradient (NDVI) is categorized in three bins (1: 0–0.3, 2: 0.3–0.5, 3: 0.5–1) and the median household income gradient is categorized in four bins (1: < 40k CA$, 2: 40k–70k CA$, 3: 70k–100k CA$, 4: > 100k CA$). There was no statistical significance detected using the Wilcoxon signed-rank test with p-value correction using the Holm procedure.

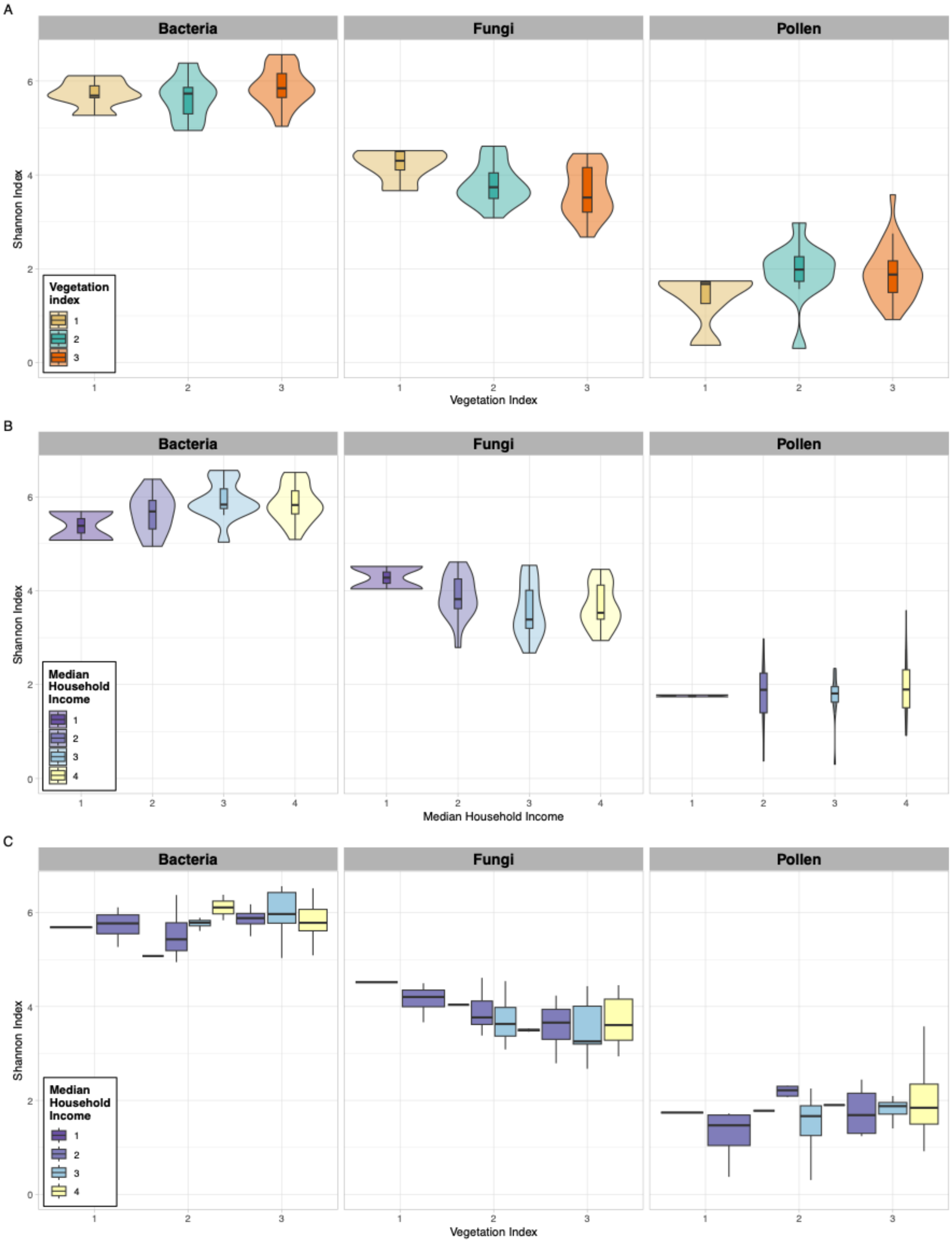

**Figure S13.** Alpha diversity across vegetation cover (NDVI) and median household income in Sherbrooke. The vegetation cover gradient (NDVI) is categorized in three bins (1: 0–0.3, 2: 0.3–0.5, 3: 0.5–1) and the median household income gradient is categorized in four bins (1: < 40k CA$, 2: 40k–70k CA$, 3: 70k–100k CA$, 4: > 100k CA$). There was no statistical significance detected using the Wilcoxon signed-rank test with p-value correction using the Holm procedure.

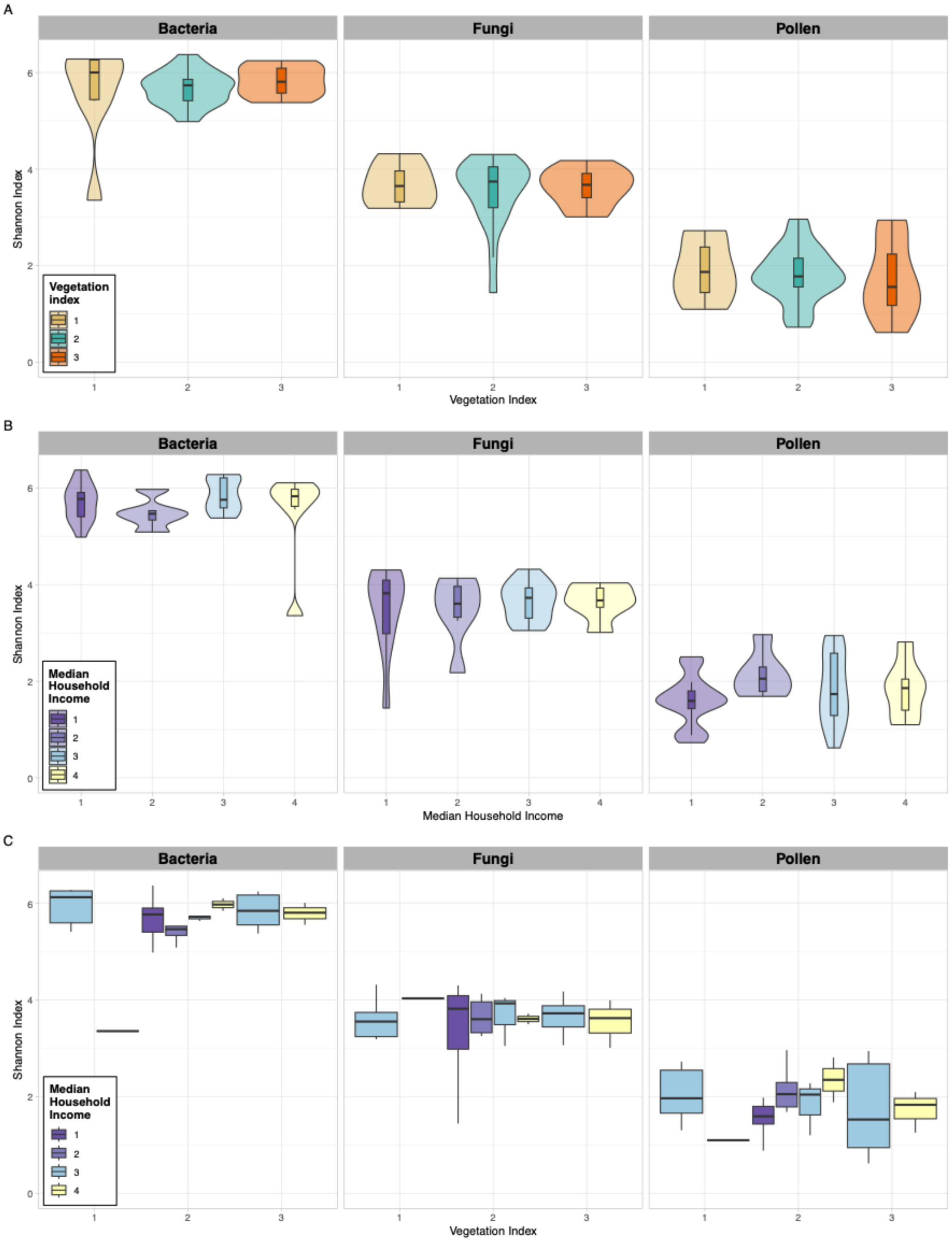

**Figure S14. Abundance of bacterial urban bioaerosols across vegetation and income gradients**. Extreme values with copy number > 5e+06 were excluded from the statistical test and plots. Values shown are log10 (mean copy number). Mean copy number is the mean of 3 qPCR replicates per sample. There was no statistical significance detected using the Wilcoxon signed-rank test with p-value correction using the Holm procedure.

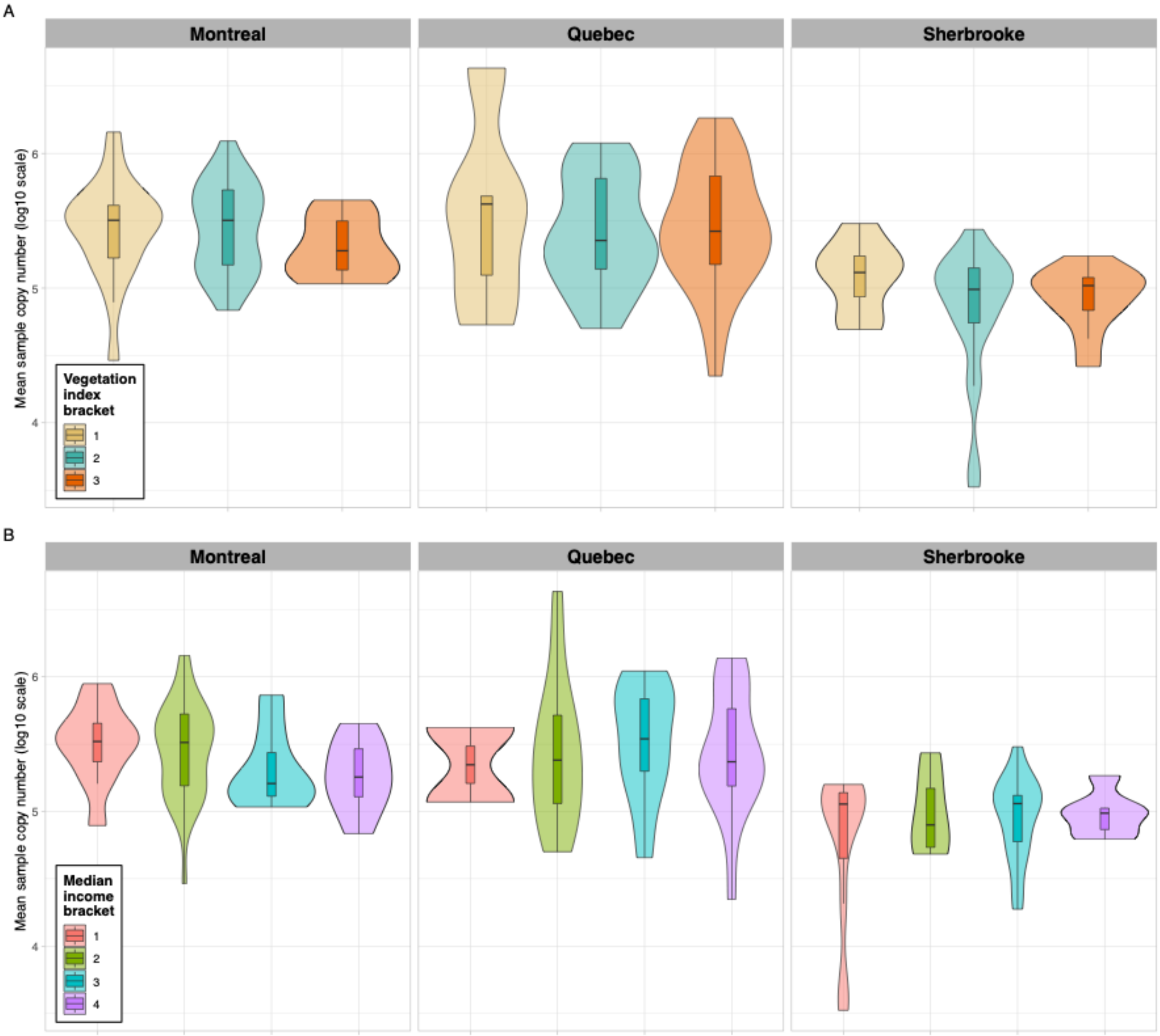

96 **Figure S15.** Sample ASV classification rates by amplicon dataset.

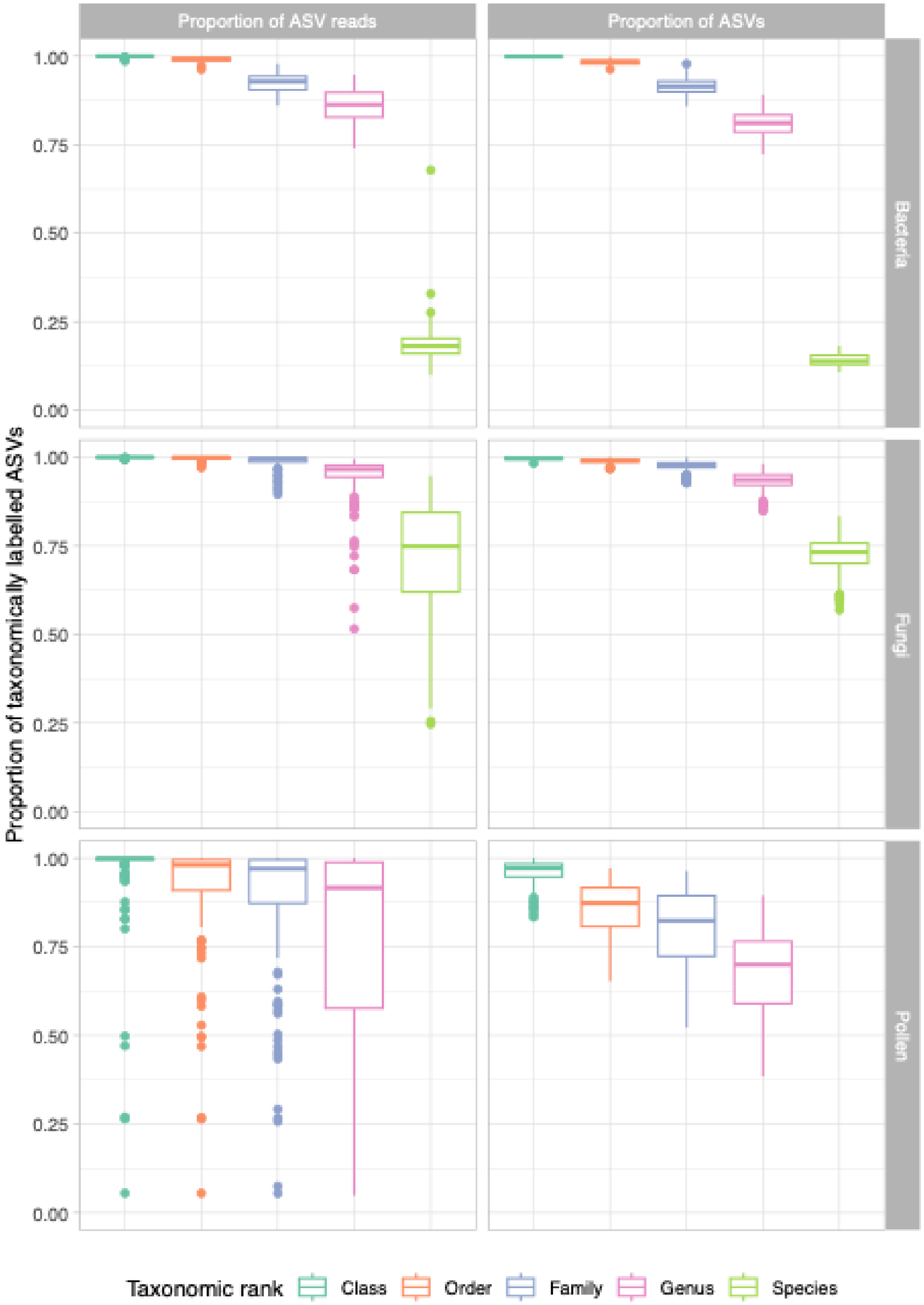


97
98
99

**Figure S16.** Mock *trnL* community sequencing. Left panel shows the relative abundance of DNA of strains from each family that was used to prepare the libraries. Right panel shows the relative sequence counts obtained based on ASVs taxonomically annotated at the family level.

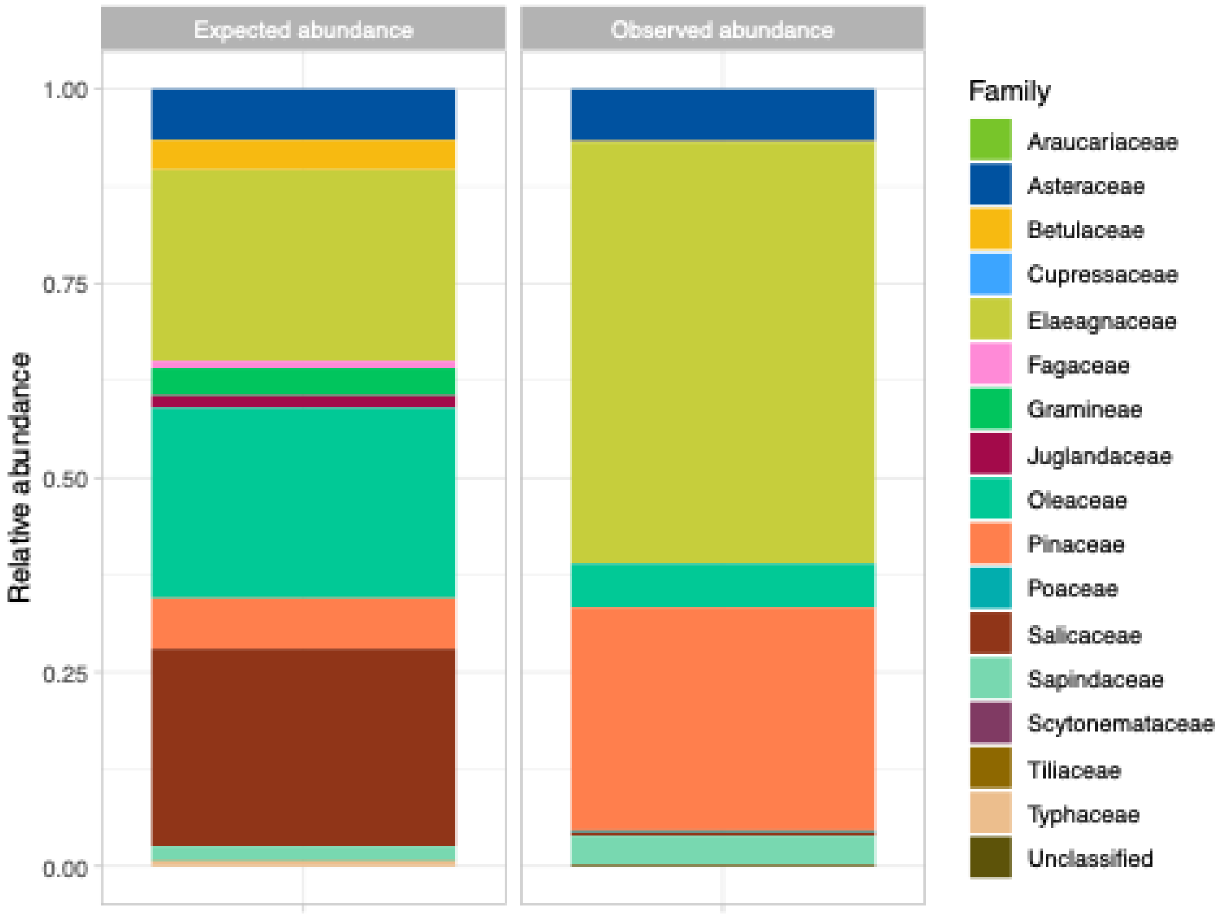

### References


1. Montreal topographic map, elevation, terrain. *Topographic maps* https://en-ca.topographic-map.com/map-mrcm14/Montreal/?center=45.57977%2C-73.57615.
2. Quebec topographic map, elevation, terrain. *Topographic maps* https://en-ca.topographic-map.com/map-tkl5k/Quebec/.
3. Sherbrooke topographic map, elevation, terrain. *Topographic maps* https://en-ca.topographic-map.com/map-k11tj/Sherbrooke/?center=45.38114%2C-71.94809.
4. Government of Canada, S. C. Population and demography statistics. https://www.statcan.gc.ca/en/subjects-start/population_and_demography (2019).
5. Canada, E. et C. climatique. Résultats de station - Données historiques - Climat - Environnement et Changement climatique Canada. https://climat.meteo.gc.ca/historical_data/search_historic_data_stations_f.html?searchType=stnProx&timeframe=1&txtRadius=25&selCity=&optProxType=park&selPark=48%7C8%7C69%7C44%7Cparc+national+Saguenay-Saint-Laurent&txtCentralLatDeg=&txtCentralLatMin=&txtCentralLatSec=&txtCentralLongDeg=&txtCentralLongMin=&txtCentralLongSec=&txtLatDecDeg=&txtLongDecDeg=&StartYear=1840&EndYear=2022&optLimit=specDate&Year=2022&Month=4&Day=5&selRowPerPage=25 (2022).
6. Aydogan, E. L., Moser, G., Müller, C., Kämpfer, P. & Glaeser, S. P. Long-term warming shifts the composition of bacterial communities in the phyllosphere of Galium album in a permanent grassland field-experiment. *Front. Microbiol.* **9**, 144 (2018).
7. Bodenhausen, N., Horton, M. W. & Bergelson, J. Bacterial communities associated with the leaves and the roots of Arabidopsis thaliana. *PLOS ONE* **8**, e56329 (2013).

8. Gardes, M. & Bruns, T. D. ITS primers with enhanced specificity for basidiomycetes - application to the identification of mycorrhizae and rusts. *Mol. Ecol. Mol. Genet. J. Wiley Online Libr.* **2**, 113–118 (1993).

9. White, T. J., Bruns, T. D., Lee, S. B. & Taylor, J. W. Amplification and direct sequencing of fungal ribosomal RNA genes for phylogenetics. in *PCR - Protocols and Applications - A Laboratory Manual* 315–322 (Academic Press, 1990).

10. Taberlet, P. *et al.* Power and limitations of the chloroplast trnL (UAA) intron for plant DNA barcoding. *Nucleic Acids Res.* **35**, e14 (2007).
